# Resilience in zebrafish embryoids

**DOI:** 10.1101/2025.06.18.660370

**Authors:** Svetlana Jovanic, Julia Eckert, Thierry Savy, Nadine Peyrieras

**Author notes:** Correspondence: N. P.

## Abstract

It is unresolved how morphogenetic cell movements and Nodal signalling are involved in the formation of the embryonic body during gastrulation. We address this question by using the zebrafish embryoid model system, which consists of cultured blastula cells lacking yolk and signalling derived from the yolk syncytial layer (YSL). Zebrafish embryoids form a cylindrical-like structure under serum-free growth culture conditions. We find that embryoids elongate without directed convergence and maintain their proliferation rate and cell mixing behaviour. Surprisingly, they undergo budding activity that has not been reported in zebrafish embryos yet. Here, the constitutively active Nodal receptor type I promotes internalisation-like gastrulation movements and induces the expression of shield genetic markers in embryoids. This work demonstrates that the absence of the yolk and signalling derived from the YSL does not affect the cell behaviour, while the constitutively active Nodal receptor type I impacts budding, internalisation movement and the emergence of the second shield.

**Summary statement:** We demonstrate that the cell behaviour during gastrulation is robust to the deficiency of yolk topology and signalling originating from the yolk syncytial layer.

## Introduction

Gastrulation is a crucial step in embryonic development in which the initial body arrangement is determined by creating the body axes (anterior-posterior, dorso-ventral) and the three germ layers (ectoderm, mesoderm and endoderm) (Gilbert and Gilbert, 2000; Holtfreter, 1944; Kimmel et al., 1995). This process is controlled by changes in gene expression, signalling pathways and cellular dynamics. Gastrulation Cell movements during gastrulation consist of epiboly, involution, ingression, convergence and extension. How these morphogenetic movements shape the embryonic body remains to be elucidated. One of the major signalling pathways involved in gastrulation is called Nodal. The Nodal pathway is involved in the specification, involution and ingression of the mesendoderm (Feldman et al., 1998; Liu et al., 2018; Sepich et al., 2005; Solnica-Krezel, 2005). However, one of the unresolved questions relates to the Nodal signalling effects on cell dynamics during axis elongation (Feldman et al., 1998; Ninomiya et al., 2004; Solnica-Krezel, 2005; Williams and Solnica-Krezel, 2020). Nodal is strongly influenced whether its pathway proteins are activated or inactivated. In the zebrafish embryo high levels of this morphogen are required at the dorsal margin to generate anterior axial mesoderm and endoderm (target genes *goosecoid* (*gsc*) and *sox32*, respectively) (Chen and Schier, 2001; Dougan et al., 2003; Gritsman et al.; Soh et al., 2020). Moreover, injection of the constitutively activated Nodal receptor type I (*acvr1ba\**) into a marginal cell at the 16-cell stage of a zebrafish embryo has been shown to induce a second dorsal organiser (shield) and give rise to the entire secondary axis in 25 % of embryos (Aoki et al., 2002; Peyriéras et al., 1998; Xu et al., 2014). The Nodal pathway is also disrupted in zebrafish lacking the maternal and zygotic one-eyed-pinhead gene (MZ*oep*) encoding the Nodal co-receptor (Gritsman et al., 1999). MZ*oep* embryos do not undergo proper gastrulation and express a shorter A-P axis (Carmany-Rampey and Schier, 2001; Montero et al., 2005; Schier et al., 1997; Strähle et al., 1997). Another effect on the Nodal pathway appears when *acvr1ba\** injected into a marginal cell at the 16-cell stage of MZ*oep* mutants. This modification restores mesoderm cells below the neuroectoderm, improves the A-P axis length and neural tube morphology (Araya et al., 2014). Taken together, these studies demonstrate that the Nodal signalling pathway is important for body axis formation and precise gastrulation. The consequences of such changes in the Nodal signalling pathway on cell movements during gastrulation needs further investigation.

Here, we investigated morphogenetic movements during gastrulation and the influence of Nodal signalling pathway using the zebrafish embryoid model (Fulton et al., 2020; Schauer et al., 2020). Embryoids are systems containing cultured zebrafish blastula cells that are capable of forming a sphere shape and then organising into cylinder-like structures. Previous studies have shown that zebrafish embryoids (explants) are proper tools for studying symmetry breaking, elongation, patterning and differentiation (Fulton et al., 2020a; Schauer et al., 2020; M. L. Williams and Solnica-Krezel, 2020). Zebrafish embryoid cells develop without the yolk. Thus, their development is not affected by the topology and biochemical activity of the yolk. Therefore, using an *in vitro* system without mechanical and geometrical boundaries of the extraembryonic tissue (yolk) has the potential to exhibit evolutionarily more conserved developmental mechanisms (Duboule, 1994; Huch et al., 2017). In addition, embryoids are half the size of a whole zebrafish embryo and are therefore more suitable for 3D+time imaging.

Although the involvement of Nodal in embryoid elongation and mesendoderm formation has been extensively studied, its influence at the single-cell level during gastrulation has not yet been reported (Schauer et al., 2020). Additionally, Recent results from this *in vitro* model also indicate the importance of elongation in maintaining an appropriate balance between BMP and Wnt signalling pathways during zebrafish gastrulation (Fulton et al., 2020a). However, how embryoids elongate at the level of cell behaviour still remains to be explored.

In this study, we first investigated cell behaviour during gastrulation in embryos and embryoids. Second, we focused on the role of Nodal signalling during gastrulation and axis elongation using the embryoid model. We found that the level of cell proliferation and mixing was similar between zebrafish embryo and embryoid. We suggest that embryoids are able to elongate without directional convergence. Moreover, we examined differences in embryoid elongation as a function of the condition of the Nodal signalling pathway. We detected tissue deformation in the form of buds that have not yet been reported in zebrafish embryos. Embryoids generated from embryos injected with *acvr1ba\** into a marginal cell at the 16-cell stage showed internalization-like movement and the development of up to two proximal extensions tips. Moreover, *acvr1ba\** injected embryoids display up to two spots of Nodal-responsive gene *gsc*. In intact embryos, this behaviour corresponds to the formation of two shields that are domains of *gsc* expression (Peyriéras et al., 1998). In summary, this study showed that zebrafish embryoids are robust to the lack of yolk topology and yolk-derived signalling during gastrulation.

## Results

### Success rate of 80% in zebrafish embryoid elongation in serum-free saline

Our study focused on the elongation of embryoids in comparison to embryos and the involvement of Nodal signalling in gastrulation of embryoids. We produced embryoids by separating the blastoderm from the yolk cell just before the midblastula transition and the formation of the three germ layers. Embryoids were cultured in 1x Ringer’s solution devoid of any growth serum including growth factors (e.g., FGF, TGF-β1) that might affect morphogenesis. Our medium reduced growth serum component-induced embryoid elongation and therefore, enabled potential observations of elongations based only on the genetically-encoded self-assembly (Turner et al., 2016). It did not affect the starting point of elongation (between 7 and 8 hpf) and its duration of several hours, consistent with previous results (Fulton et al., 2020b; Schauer et al., 2020b) (Fig. 1).Using our minimal culture conditions, we obtained a success rate of more than 80% in wt embryoid elongation (148 embryoids from 14 independent experiments), showing a cylindrical-like structure (Fig. 1B).

**Fig. 1.**
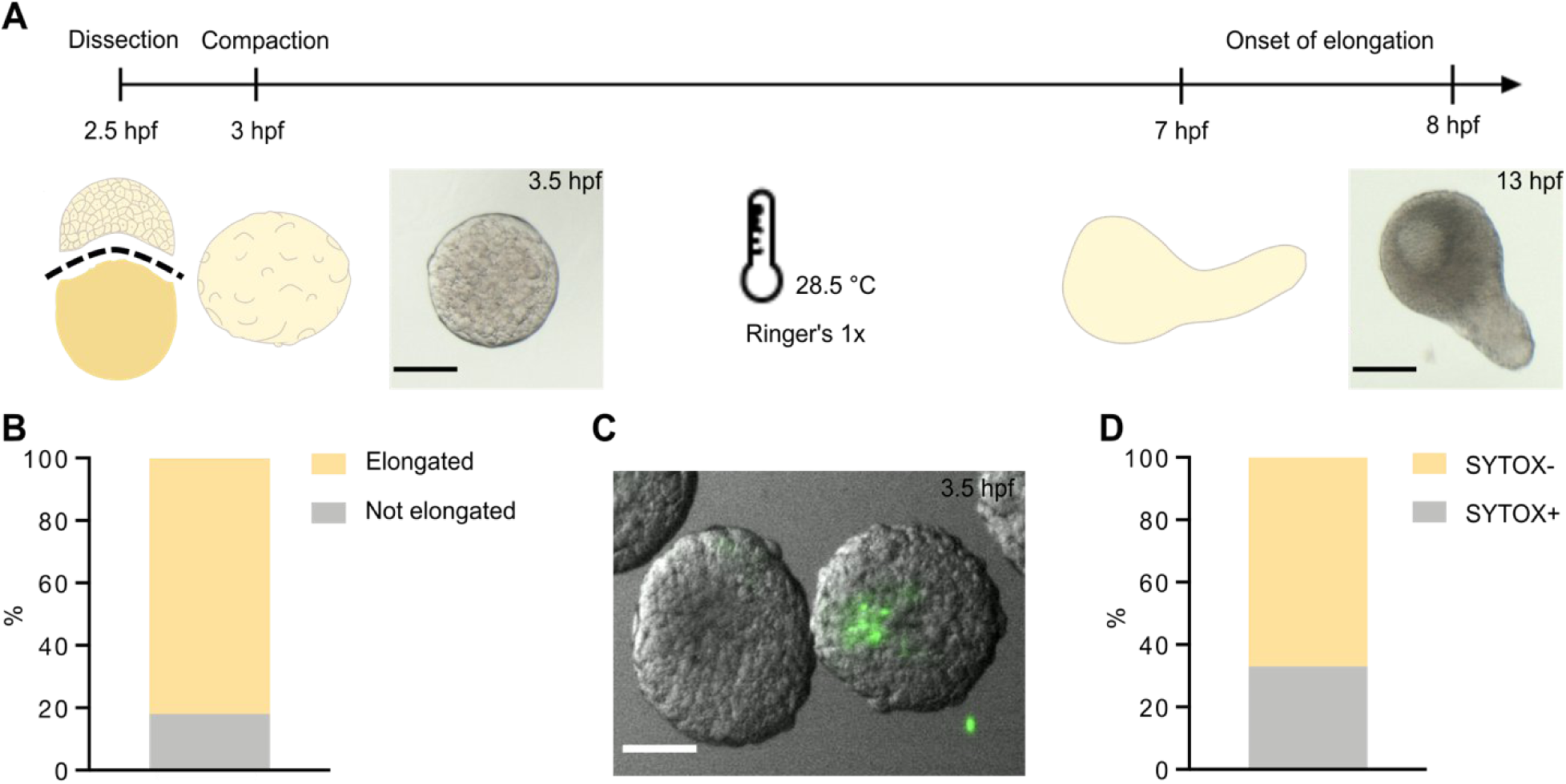
Elongation of embryoids formed from 2.5 hpf blastulae. (**A**) Protocol for embryoids and their observation; mechanical separation of the blastoderm from the yolk cell at 2.5 hpf, embryos culture in 1x Ringer’s solution at 28.5°C, compaction within the next 30 minutes (bright field image), elongation starts between 7 and 8 hpf, observation until 13 hpf (bright field image). (**B**) 80% of the embryoids elongated (N=14, n=148), (**C**) SYTOX injection into the yolk (64-cell to 128-cell stage), embryoids imaged at 3.5 hpf, (**D**) 67% of embryoids (N=1, n=18) are devoid of SYTOX staining. Scale bars: 200 µm.

In addition, we investigated the presence of yolk and yolk nuclei in our embryoids to study the Nodal signalling activity triggered by the maternal Ndr1 and maternal beta-catenin pathway (Fig. 1D). The yolk was fully eliminated in 67% of the embryoids.

By injecting SYTOX Green fluorescent dye into the yolk (64-128 cell stage) before the dissection, we demonstrated that embryoids were almost entirely devoid of yolk. In only 33% of our embryoids at stage 3 hpf, we observed traces of yolk nuclei (Fig. 1C).

## Embryoids elongate without directed convergence

To test the mechanism of convergence and extension in embryoids, we analysed the movements of digitally detected cell centres relative to an arbitrarily selected midline. We selected four regions, left (red), right (blue), top (white) and bottom (yellow) to compare cell movements in the embryo and embryoid at different observation angles. Selected cell trajectories were followed between an observation time window of 6.5 and 12.5 hpf. These trajectories were represented in rotation angles of 0°, -45° and +45° to obtain significant recognition of convergence and extension (Fig. 2A).

**Fig. 2.**
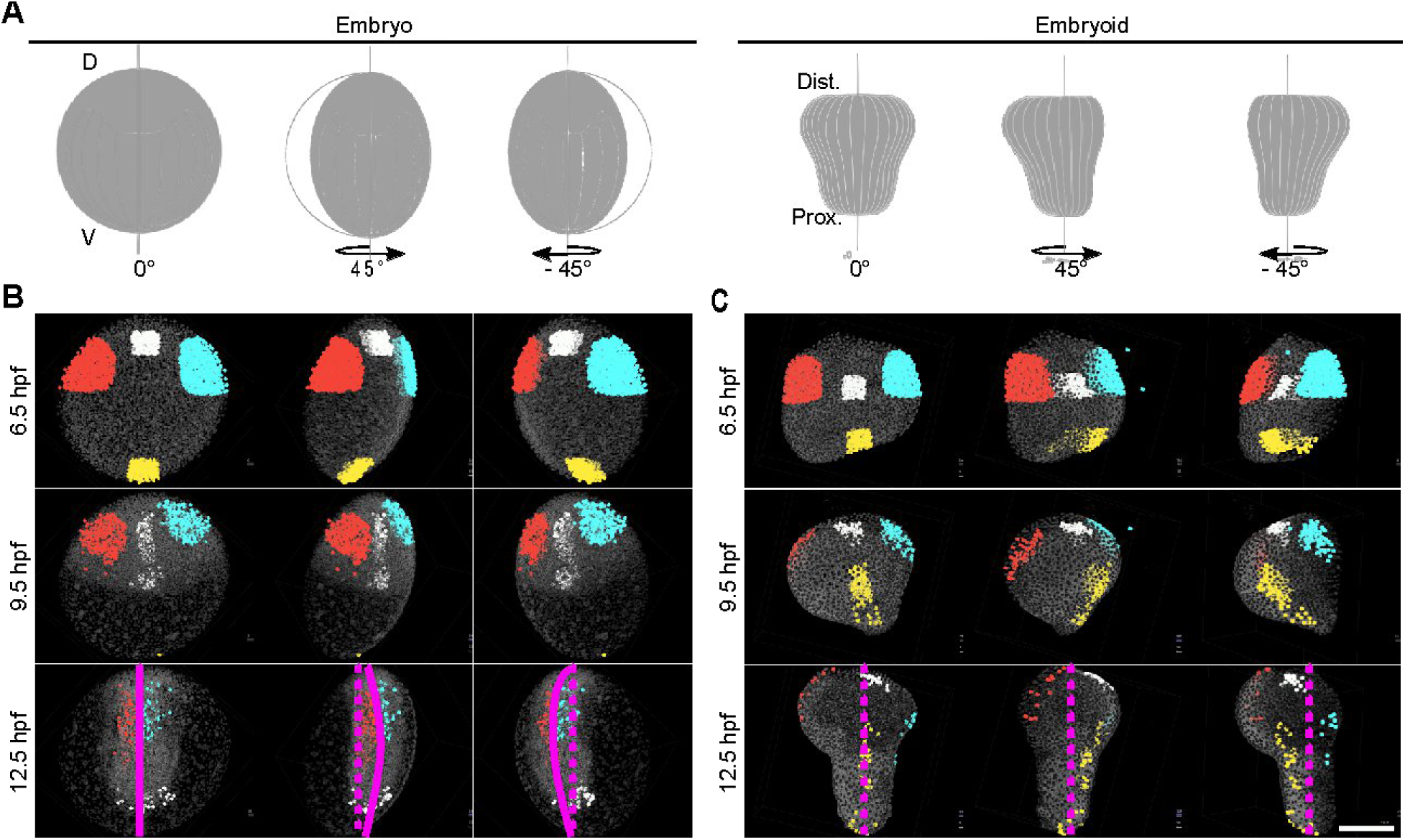
Comparison of embryo and embryoid cell behaviour. (**A**) Schematics represents starting positions of the embryo (left) and embryoid (right) presented from the top view at 6.5 hpf and rotated for 0°, 45° and -45° to the y-axis. D, dorsal, V, ventral (embryo), Prox., proximal, Dist., distal (embryoid). Snapshots of the embryo (**B**) and embryoid (**C**) maximum projections ubiquitously expressing H2B-mCherry (grey) in nuclei superimposed to digitally selected nuclei centres coloured in red, blue, white and yellow. Snapshots corresponded to 6.5, 9.5 and 12.5 hpf. Embryo and embryoid are oriented as presented on schematics. The real midline is shown as a magenta line and the arbitrary midline is given as dashed magenta line, bottom row. Scale bar: 200 µm.

In the embryo, convergence and extension were observed from 8 - 12.5 hpf (Fig. 2B). Convergence was indicated by the movement of cells (red and blue selections) toward the midline, followed by a downwards movement along this midline (extension), as seen at 0°. To prove the existence of only one midline in the embryo on which convergence occurs, we rotated the sample by +/-45° and selected a new arbitrary midline (dotted line). The selected red and blue cells did not follow the convergence movement along this line. Cells were localised at a certain distance from the arbitrary midline and moved along the curved real midline selected at 0°. This result showed the existence of only one midline in the embryo along which convergence occurs. Once the cells reached the convergence midline, elongation took place, as can be seen for the white selected cells at 9.5 hpf and later for the red and blue selected cells at 12.5 hpf. Cells coloured in yellow moved out of field of view (FOV) to posterior regions. Surprisingly, we did not observe convergence movements of cells in embryoids compared to the embryos (Fig. 1C, Movie 1). The red and blue selected cells did not reach the midline over time, which was not observed at 0° nor at 45° positions. These cells migrated downward and remained along the surface of the lateral sides of the embryoid. Only extension was observed. The cells belonging to the region of the embryoid marked in white did not follow the extension, as represented in the embryo, but sustained along the surface on the top part of the embryoid. The yellow-marked cell populations were located at the proximal part of the embryoid, elongated in the same direction over time, and narrowed moderately due to the extension process.

**Movie 1.**
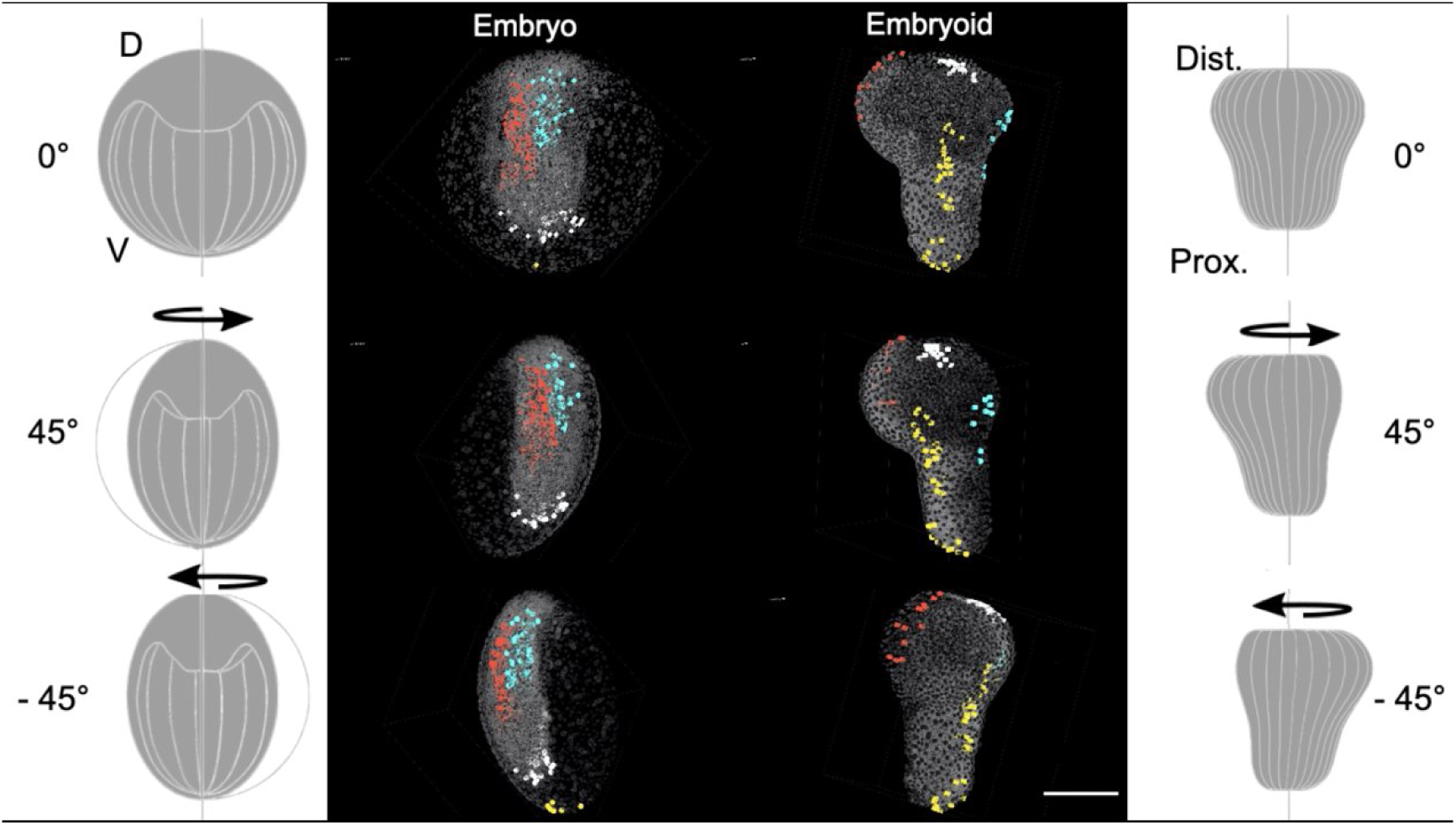
Assessment of convergences towards a midline. Maximum intensity projections for embryo and embryoid raw data superimposed with four selected cell centre populations. Illustrations on the left and right present embryo (left) and embryoid (right) rotated for 0°, 45° and -45°. Four populations of nuclei centre selections (coloured in red, blue, white and yellow) forward tracked from 6.5 to 12.5 hpf in control embryo and embryoid. Samples ubiquitously expressed mCherry-H2B in nuclei (grey). D, dorsal, V, ventral, Prox., proximal, Dist., distal. Data acquisition parameters are given in Table S3. 2. Scale bar: 200 µm.

Our results indicated that constraints and signalling derived from extraembryonic tissues in the absence of yolk. Embryoids follow a different path in their development. The direction of collective cell movements causes only an extension compared to embryos, but not convergence.

Considering that directed convergence on the midline has not been identified in embryoids, we further deciphered this observation by examining the spatial organisation of krox20 (marks hindbrain rhombomeres 3 and 5) in this model (Movie 2). In the embryo, krox20 is expressed at the convergence region, which we have indicated as the midline in a rounded, arch-like shape. In embryoids, krox20 emerges as only one half of a round arch-like shape. The absence of the second part of krox20 in embryoids further implied a convergence-midline deficiency. Therefore, we can report a broken symmetry of krox20 in embryoids given in embryos according to the midline.

**Movie 2.**
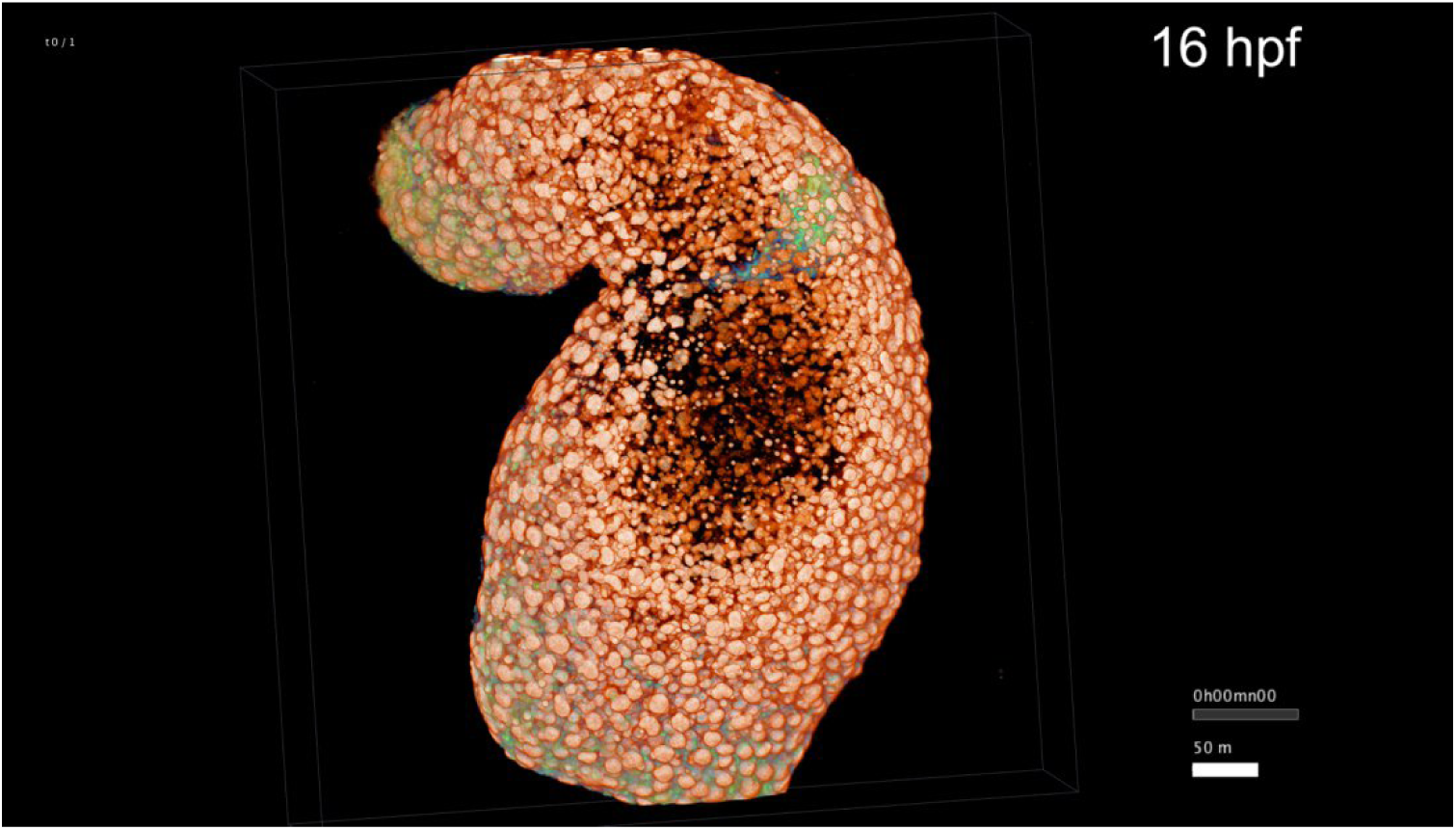
*Krox20* reporter spatial expression pattern in embryoid (16 hpf). Embryoid expressed ubiquitously H2B-mCherry in nuclei (orange) and *krox20* driven EGFP membrane expression (green). 3D representation of *krox20* spatial expression in embryoid indicating its position in one quarter of the embryoid width. Data acquisition parameters are given in Table S3. 3. Samples oriented from proximal (top) to distal (bottom). Scale bar: 50 µm.

### Cells in embryoids maintain their proliferation rate and cell mixing behaviour

Next, we wanted to determine whether cells in embryoids in the absence of the yolk behave in the same way in terms of proliferation and movements as in the embryo during symmetry breaking and elongation. We reconstructed the cell lineages of embryoids and zebrafish embryos between a few minutes before the start of gastrulation (5 and 5.3 hpf) to around 12 hpf. Based on the duration between two divisions belonging to the same lineage, we found that embryos and embryoids had a similar cell cycle duration (p=0.19) (Fig. 3A,C). Furthermore, we calculated how much cells belonging to the central and marginal regions mix. Embryo and embryoid did not show significant differences (p=0.17) (Fig. 3B,D). Taken together, our measurements indicated that the cell behaviour in terms of proliferation rate and cell mixing is robust to the absence of the yolk topology and YSL-derived signalling.

**Fig. 3.**
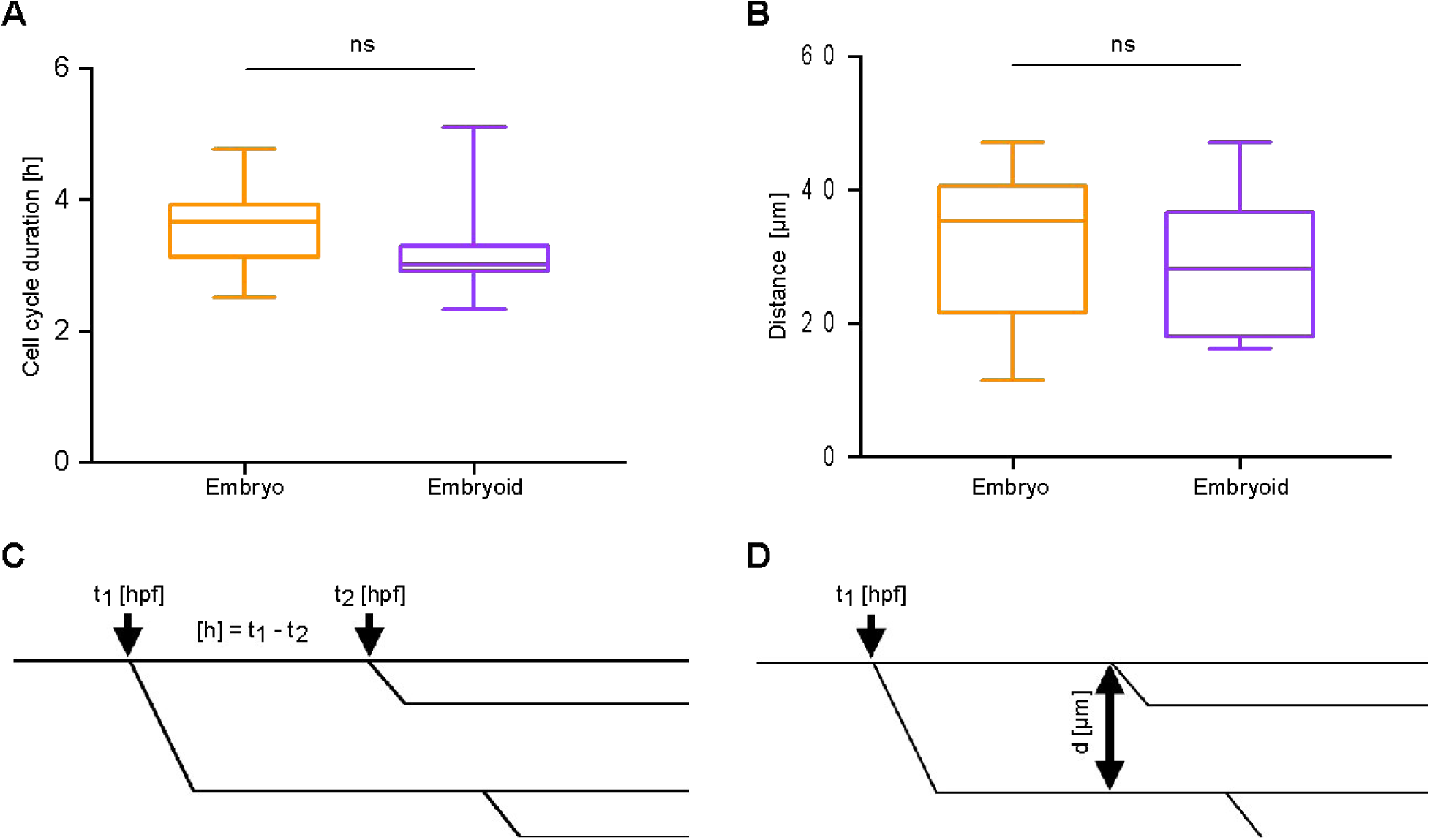
Comparison of embryo and embryoid in terms of proliferation rate and cell mixing. The difference in wt embryo and embryoid in terms of (**A**) proliferation (1 embryo 17 cells, 1 embryoid 12 cells) and (**B**) cell mixing (1 embryo 10 cells, 1 embryoid 21 cells) from 5 to 12 hpf and 5.3 to 12.2 hpf, respectively. Groups compared using unpaired t-test. Data are shown as boxes that extend until the 25th and 75th percentiles with the middle line presenting the median, error bars are 1-99 percentiles. ns, not significant, p=0.1865 and p=0.1657, respectively. (**C**) Cell cycle duration is given as a difference between time point one of the daughter’s divides (t_2_) and time point mother divides (t_1_), bottom left. (**D**) Mixing is assessed by measuring the distance in µm between sister cells when one of them divides, bottom right.

## Modulation in Nodal signalling activity affects embryoid elongation

To determine whether Nodal signalling is involved in embryoid elongation the following embryos were generated: wild type (wt), wild type injected with constitutively active receptor *acvr1ba*\* (wt+*acvr1ba*\*), Nodal-deficient mutants (MZ*oep*) and MZ*oep* mutants injected with *acvr1ba*\* (MZ*oep*+*acvr1ba*\*) (Fig. S1, Table S1, Movies 3, 5). Using time lapse bright field imaging, we observed that wild type embryoids form one proximal extension tip at around 8 hpf. Surprisingly and in contrast, 18% of wt+*acvr1ba*\* formed up to two dominant extensions (Table S1). Despite the absence of *oep* co-receptor in MZ*oep* mutants and MZ*oep*+*acvr1ba*\*, some embryoids succeed to elongate (Fig. 4A).

**Fig. 4.**
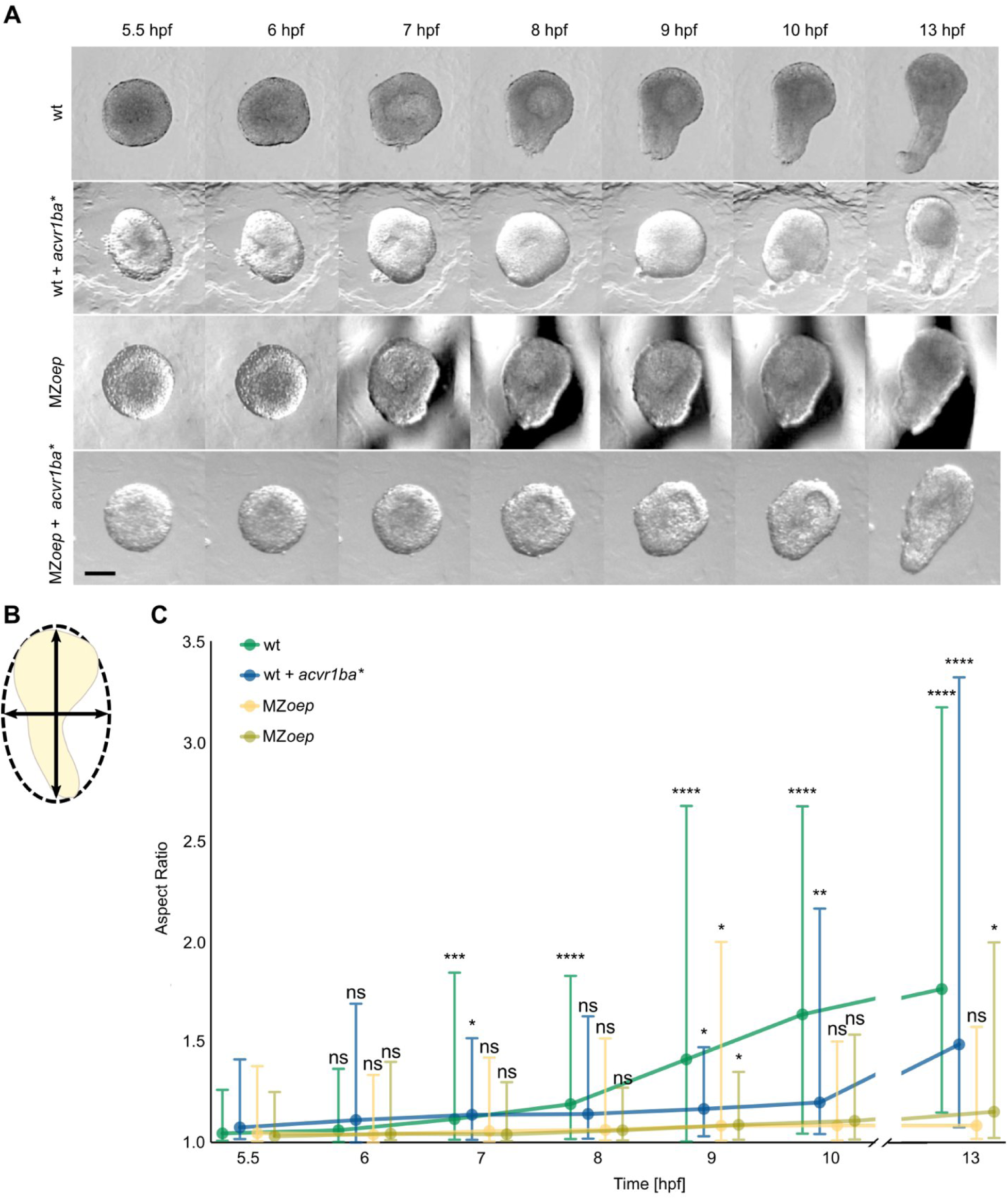
Quantification of shape change depending on Nodal signalling. (**A**) Bright field two-dimensional images represent four embryoid groups (wt, wt+*acvr1ba*\*, MZ*oep* and MZ*oep*+*acvr1ba*\*) observed from 5.5 to 10 hpf and at 13 hpf. Images represent examples with highest level of aspect ratio and in case of wt+*acvr1ba*\* with two proximal tips. Variation in morphology is assessed by aspect ratio. Embryoids oriented from distal (top) to proximal (bottom). (**B**) Schematics depicts aspect ratio as a proportion between major and minor diameter (arrowed lines), orthogonal to each other, of ellipsoid fit (dotted line). (**C**) Aspect ratio over 7.5 h for four types of embryoids (wt, N=3, n=56; wt+*acvr1ba*\*, N=4, n=36; MZ*oep*, N=4, n=36; MZ*oep*+*acvr1ba*\*, N=3, n=28). Groups compared with Mixed-effects analysis multiple comparisons 2-way ANOVA test. Data are shown as median dots; error bars represent the range. ns, not significant, *p<0.05, **p<0.0, ***p<0.001, ****p<0.0001. Scale bar: 200 µm.

**Fig. S1.**
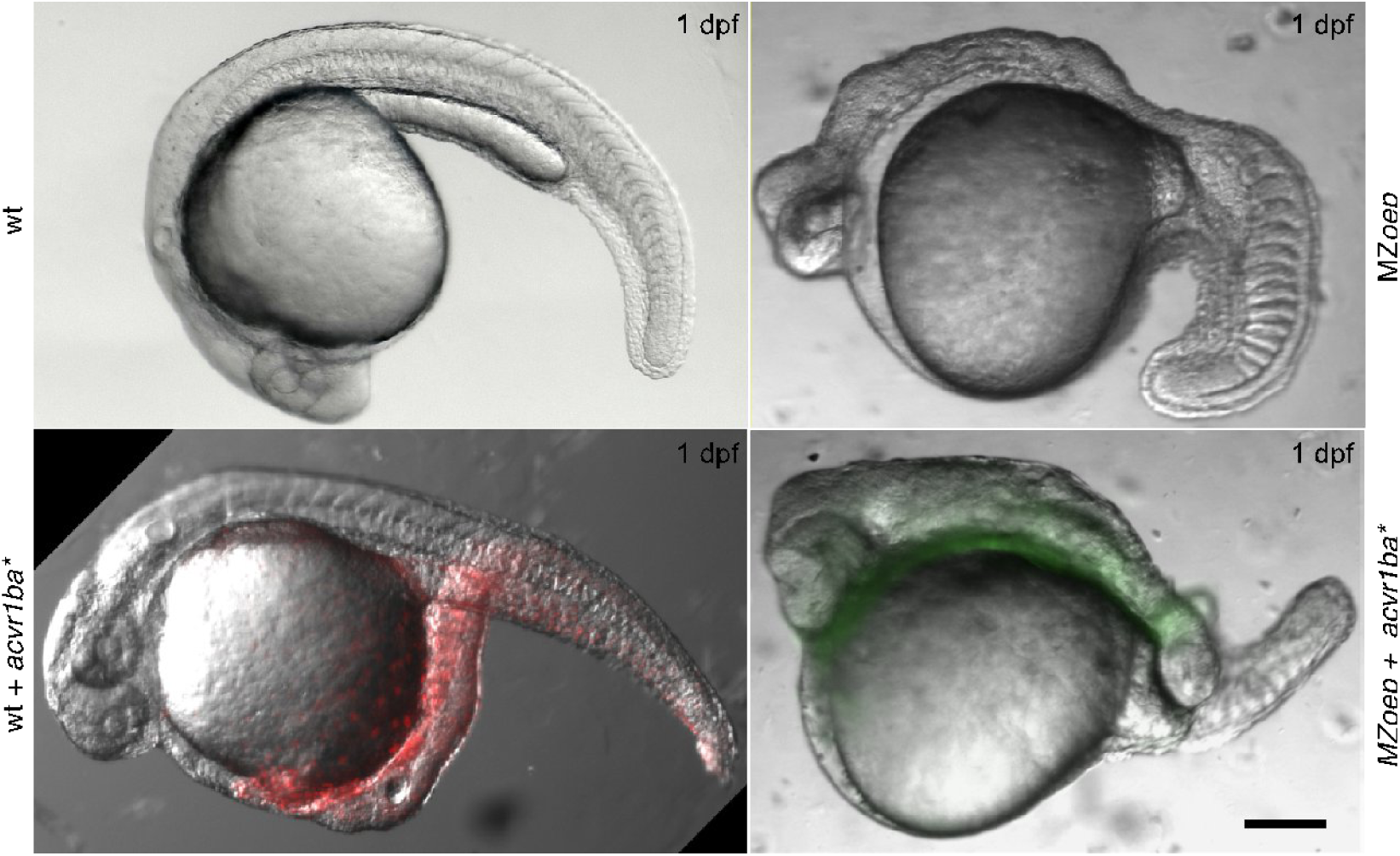
Phenotypic classes of embryos (1 dpf) depending on Nodal activity. Related to. Fig. 4. Top left, the image represents the wt embryo. Bottom left, image shows an overlap between bright field wt+*acvr1ba*\* embryo and injected progeny of 1/16 cell stage injection of mRNA encoding *acvr1ba*\* together with *H2B-mCherry* (red). Top right depicts MZ*oep* embryo. MZ*oep*+*acvr1ba*\* represents bright field and fluorescence signal originating from 1/16 cell injection progeny of *acvr1ba*\* together with *Hras-EGFP*, bottom right. Scale bar: 200 µm.

**Movie 3.**
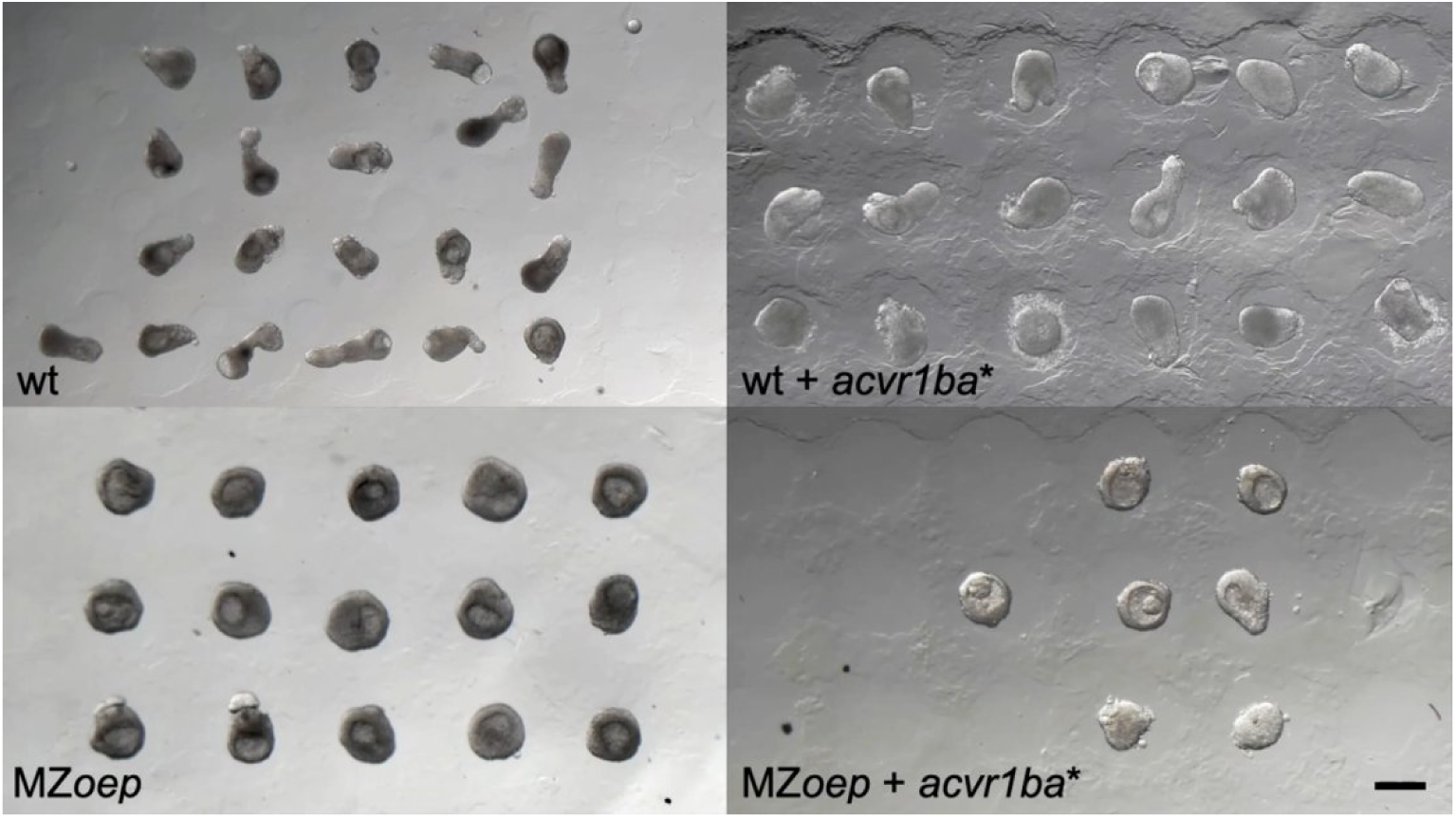
Bright field 2D+time data for embryoids depending on Nodal activity. Embryoids belonging to the following Nodal signalling pathway conditions: wt, wt+*acvr1ba\**, MZ*oep* and MZ*oep+acvr1ba*.* Observation time from 5.5-13 hpf. Data acquisition parameters are given in Table S3. 5. Scale bar: 400 µm.

**Movie 4.**
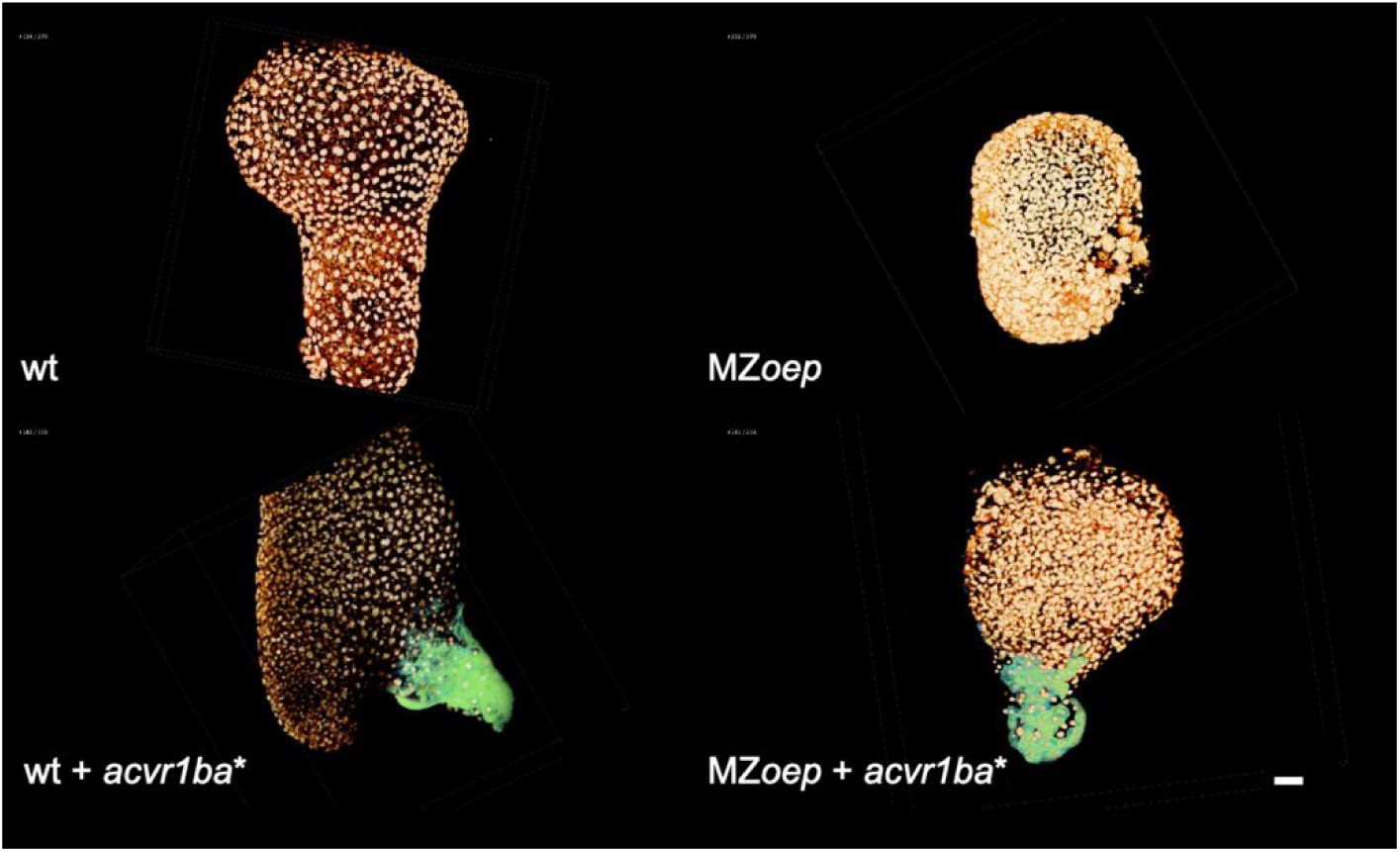
3D+time data for embryoids depending on Nodal activity. Embryoids belong to the following Nodal signalling pathway conditions: wt, wt+*acvr1ba*\*, MZ*oep* and MZ*oep*+*acvr1ba*\* developing from 5.5 to 13 hpf. Data acquisition parameters are given in Table S3. 2. Embryoids ubiquitously expressing mCherry-H2B in nuclei (orange) and Hras-EGFP with *acvr1ba*\* in the progeny of one cell on the margin microinjected at 16 cell stage (green). Samples oriented from distal (top) to proximal (bottom). Scale bars: 50 µm.

**Movie 5.**
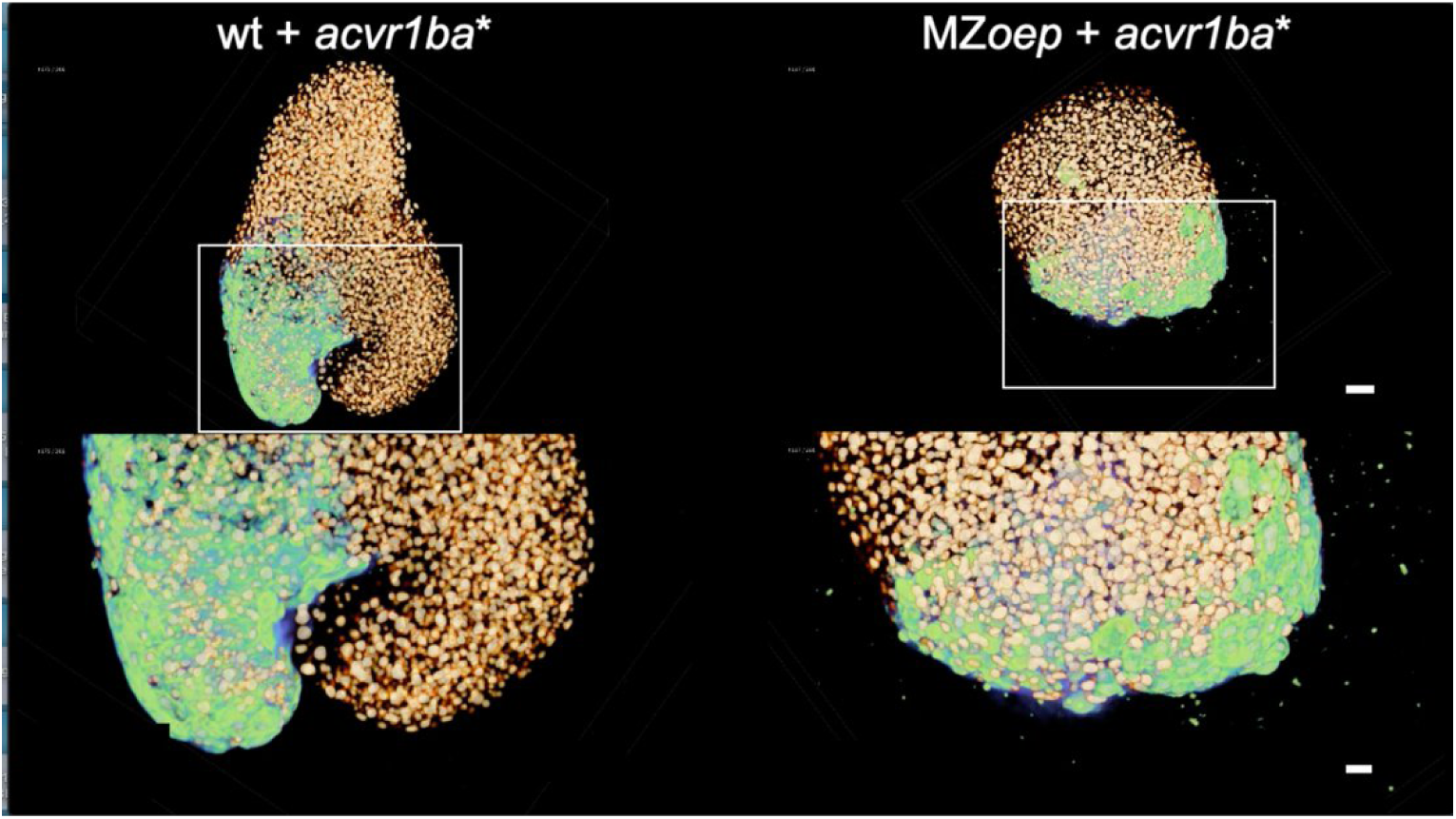
Internalisation-like movements in wt and MZ*oep* mutant embryoids upon 1/16 cell injection of *acvr1ba*\*. mCherry-H2B ubiquitously expressed in nuclei (red) and Hras-EGFP with *acvr1ba*\* in the progeny of one cell on the margin microinjected at 16 cell stage (green). Zoom-in segments, white rectangles, depicted collective cell movements appeared outside of the embryoid, synchronised return to the embryoid and emergence of two proximal extension tips. Samples oriented from distal (top) to proximal (bottom). Scale bars: 50 µm.

**Table S1.**
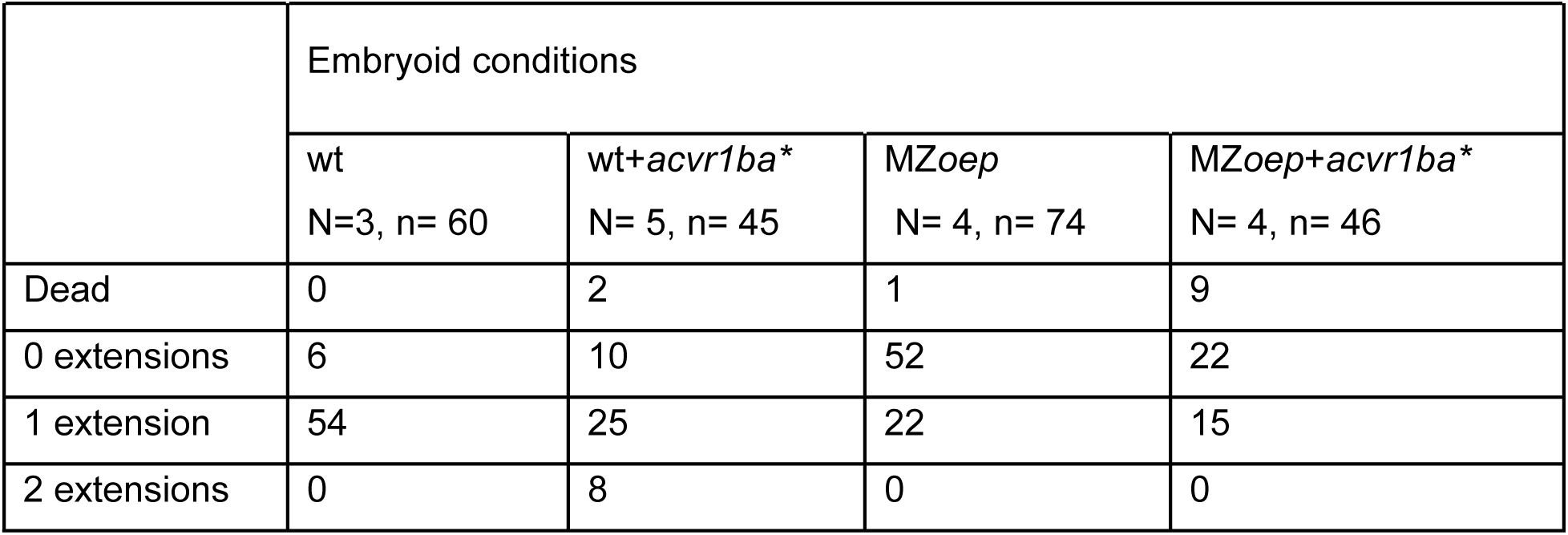
Number of proximal extensions depending on Nodal activity (13 hpf).

To compare the extension development over time (5.5 to 13 hpf) between the four embryoid conditions, we calculated the elongation as the aspect ratio (Figs 4B, S1). At 8 hpf, wild type embryoids started to extend along the major axis and shrink along the minor axis, which is known to be the first morphogenetic symmetry breaking moment in zebrafish (Keller et al., 2008). Wild type embryoids showed the most significant change in aspect ratio values over time compared to the other three conditions. Surprisingly, Nodal gain of function (wt+*acvr1ba*\*) embryoids showed smaller changes in aspect ratio values than in wt. In contrast to what has been reported (Schauer et al., 2020b; Williams and Solnica-Krezel, 2020), some embryoids (29.7%) obtained from MZ*oep* eggs achieved to extend. MZ*oep* embryoids with the addition of the constitutively active *acvr1ba*\* receptor showed a greater significant change in elongation than in MZ*oep* samples (Fig. 4C).

## Zebrafish embryoids budding activity coincides with the timing of gastrulation onset

We next focused on the budding activity of embryoids. We observed an increase in budding activity at the onset of gastrulation of embryoids that was not reported in gastrulating zebrafish embryos. During budding morphogenesis, cells collectively expand on the periphery and fold inward (Wang et al., 2021). Budding exhibits from the embryoid for several hours, starting approximately 2 hours after embryo dissection obtained at 2.5 hpf. In wt and wt+*acvr1ba\** embryoids from 7 to 10 hpf, only one of the buds took over and elongated. Embryoids belonging to MZ*oep* and MZ*oep*+*acvr1ba\** stopped the budding activity between 8 to 10 hpf and 9 to 10 hpf, respectively (Fig. 5A).

**Fig. 5.**
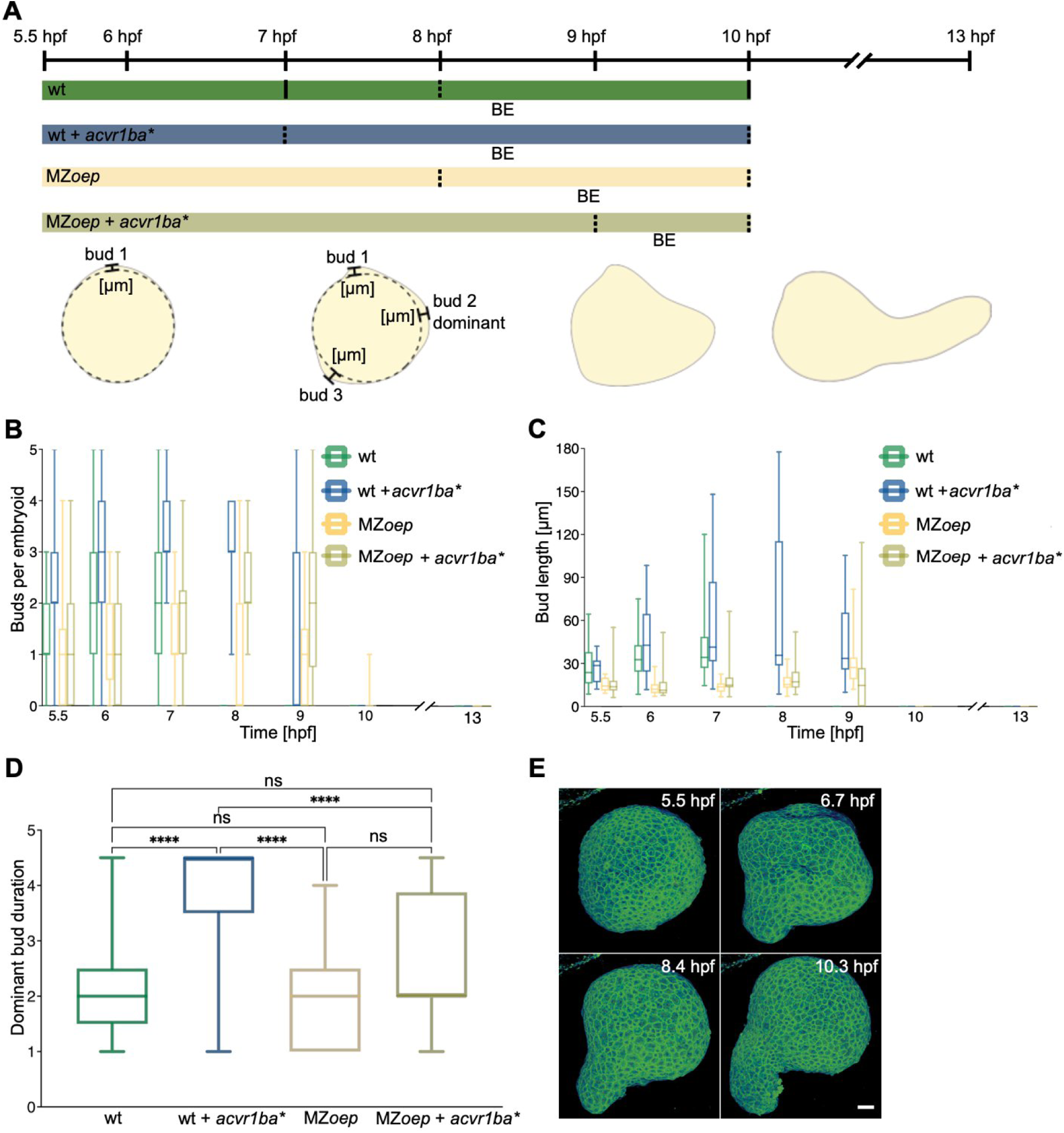
Response of embryoids to Nodal activity in terms of bud number, bud length and bud persistence. (**A**) Schematics depicts time from 5.5 hpf until budding activity stops (dotted vertical lines) for four embryoid types. The drawing represents buds located to follow their number and to measure their length. (**B**) Graph shows number of buds in single embryoid over time for four embryoid conditions (wt, N=3, n=56; wt+*acvr1ba*\*, N=4, n=31; MZ*oep*, N=3, n=49; MZ*oep*+*acvr1ba*\*, N=3, n=30). (**C**) Difference in bud length over time based on embryoid condition (wt, N=1, n=14; wt+*acvr1ba*\*, N=3, n=14; MZ*oep*, N=1, n=15; MZ*oep*+*acvr1ba*\*, N=3, n=28). (**D**) Quantification of dominant bud duration measured from 5.5 hpf until time point related bud elongates in four embryoid states (wt, N=3, n=42; wt+*acvr1ba*\*, N=5, n=29; MZ*oep*, N=2, n=17; MZ*oep*+*acvr1ba*\*, N=2, n=16). Groups compared with multiple comparison 2-way ANOVA test. Data shown as boxes that extend until the 25th and 75th percentiles with the middle line presenting median, error bars are 1-99 percentiles. ns, not significant, ****p<0.0001. (**E**) Selected maximum intensity projections represent budding activity in the control embryoid. Data acquired in 3D from 5.5-13 hpf. Buds formed depicted at time point 6.7 hpf. Embryoid derived from embryo that is ubiquitously expressing EGFP-Hras in membrane. BE, budding ends. Scale bar: 50 µm.

Next, we investigated whether different Nodal signalling conditions induce changes in the embryoid budding activity. To do so, we measured the number of buds and their length in embryoids for the period from 5.5 to 10 hpf and at time point 13 hpf, concerning four embryoids prepared from embryos with distinctive Nodal signalling status (Fig. 5B). At 5.5 hpf, we observed zero, one or two buds over the following 2.5 hours. Afterwards, the budding was replaced by one or two proximal extensions. In Nodal gain of function embryoids (wt+*acvr1ba\**), we detected at least one or more buds compared to all other embryoid conditions. MZ*oep* embryoids overexpressing *acvr1ba\** (MZ*oep*+*acvr1ba*)* revealed a larger number of buds in later stages of budding than in the MZ*oep* group. Moreover, our results showed that wt+*acvr1ba\** embryoids created longer buds compared to the other three conditions (****p<0.0001) (Fig. 5C).

Furthermore, we aimed to reveal whether the dominant bud, the bud that elongates, is the one that has the longest lifetime and whether the lifetime of the dominant bud changes corresponding to the Nodal signalling condition (Fig. 5D). To analyse the duration of the dominant bud, we measured the time between 5.5 hpf until the end of budding (BE) for each embryoid of the four embryoid states. Our results showed that the bud with the maximum lifetime was not always dominant in all four conditions. However, the dominant bud lifetime in Nodal overexpressed embryoids lasted for 2.5 h, which was longer than in the other three embryoid conditions (****p<0.0001).

It has been previously reported that zebrafish embryoids were characterised with the appearance of the cavity on the opposite side to the extension tip (Schauer et al., 2020). We observed the largest distance between the dominant bud and the cavity at the early gastrulation stage (5.5 hpf) (Fig. S2). Distance measurements were obtained for four embryoid conditions. In all conditions, the distance of the dominant buds to the cavity was reduced. Thus, in wt embryoids, 89% of the dominant buds had the largest distance to the cavity. In wt+*acvr1ba\**, MZ*oep* and MZ*oep*+*acvr1ba\** embryoids, the percentage of dominant buds decreased.

**Fig. S2.**
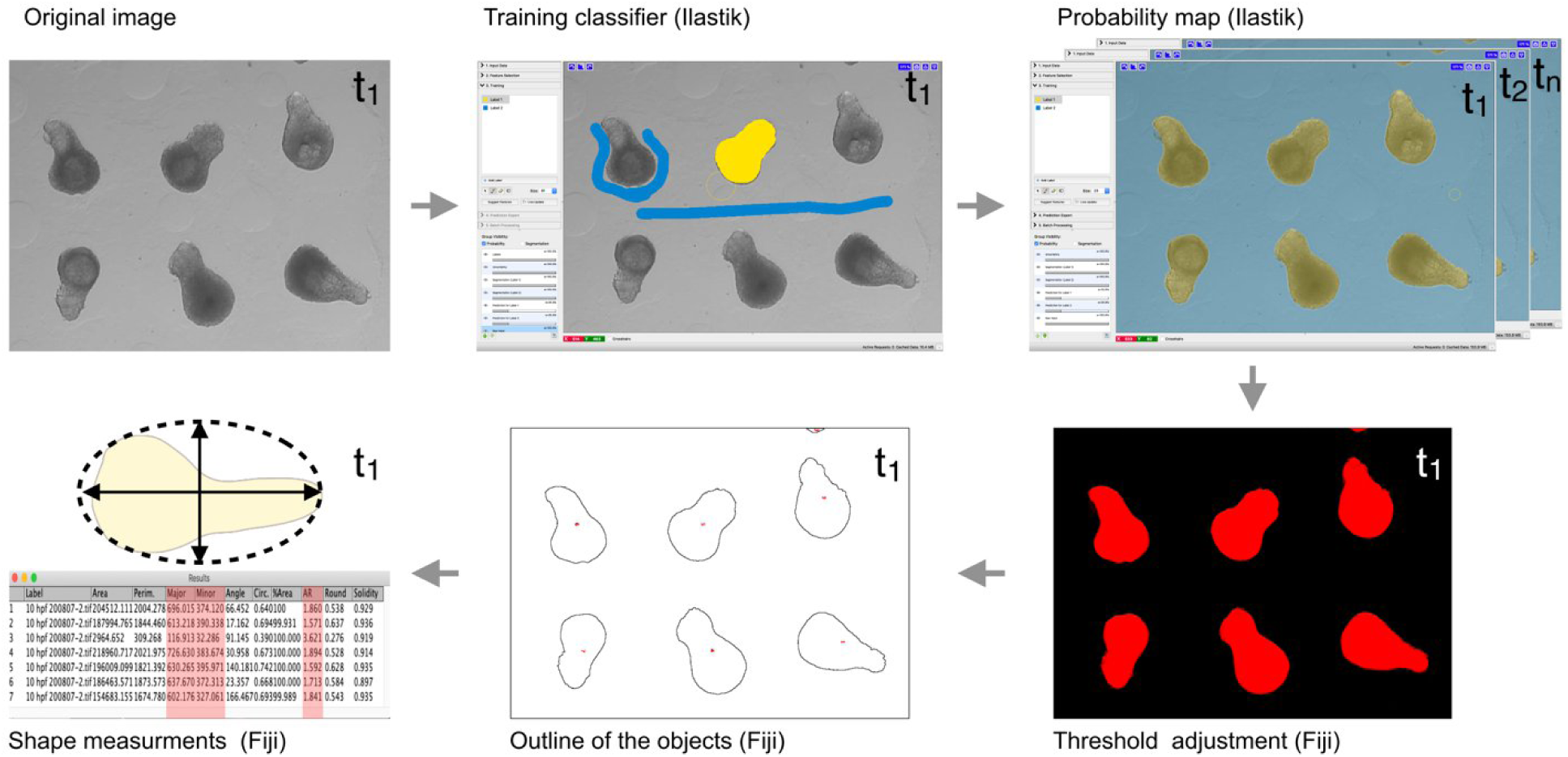
Measurement of the elongation. Related to. Fig. 4. Bright filed original images extracted from the time lapse videos for each 1 h of embryoid development. To segment embryoid shape Ilastik software (Berg et al., 2019) was used. Training with one of the images was done by Pixel Classification Ilastik workflow. After the classifier is trained, it is possible to apply results on the batch of other images and generate probability maps. Probability maps were analysed in Fiji by using threshold adjustment, analyse particles option to generate outline of embryoids and finally to calculate Aspect Ratio (AR = Major axis/Minor axis) for the best fit to ellipse.

## Expression of *acvr1ba\** in embryoids promotes internalisation-like gastrulation movements

Previous findings have shown that overexpression of Nodal signalling by *acvr1ba\** in naïve transplanted cells can lead to internalisation behaviour of cells (David and Rosa, 2001). Furthermore, it has been reported that wt embryoids do not undergo internalization (Schauer et al., 2020). Therefore, we asked whether internalisation can be induced in wt and MZ*oep* embryoids after *acvr1ba\** injection at the 16-cell stage prior to yolk dissection. Larger tissue deformations were observed from the 3D+time data with ubiquitously labelled nuclei, using reporter (*H2B-mCherry*) in embryoids made of 1/16 cell embryos co-injected with *acvr1ba* and Hras-H2B* in wt and MZ*oep* phenotypes. The movement from 6.7 to 10.5 hpf resembled morphogenetic internalisation-like movement in zebrafish embryos. Moreover, after the internalisation-like movement the second proximal tip was observed from 10.5 to 13 hpf in 18% of wt+*acvr1ba\** specimen. MZ*oep*+*acvr1ba\** express two much shorter proximal tips compared to wt+*acvr1ba\** embryoids (Fig. 6, Movie 5).

**Fig. 6.**
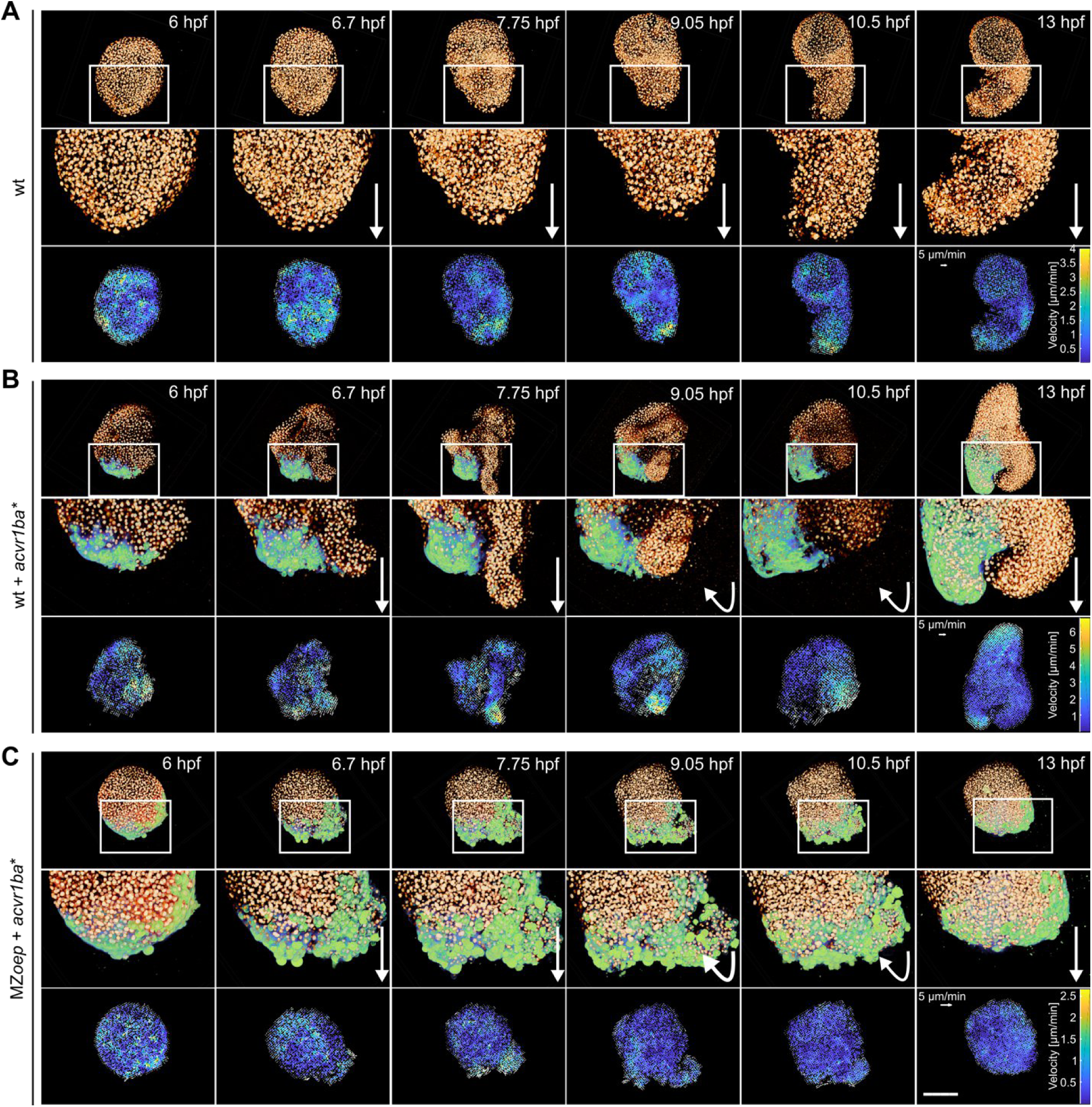
Internalisation-like movements appeared upon *acvr1ba*\* injection in wt and MZ*oep* embryoids. Selected maximum intensity projections of embryoids starting from 6 to 13 hpf of (**A**) wt expressing ubiquitously H2B-mCherry (yellow) in nuclei, (**B**) wt + *acvr1ba*\* and (**C**) MZ*oep* + *acvr1ba*\* expressing ubiquitously H2B-mCherry (yellow) in nuclei and *Hras-EGFP* with *acvr1ba*\* (green) in the progeny of one cell on the margin microinjected at 16 cell stage. Zoom-in segments, white rectangles, followed with white arrows represented collective cell movements toward the distal part of the wt embryoid; white arrowheads in wt + *acvr1ba*\* and MZ*oep* + *acvr1ba*\* represented collective cell movements to outside (6.7 and 7.75 hpf), synchronised return to the embryoid (9.05 and 10.5 hpf) and the emergence of two extensions (13 hpf) in which one or both belong to the progeny of 1/16 cell injection of *acvr1ba*\* and *Hras-EGFP*. The bottom row in each embryoid condition (wt, wt + *acvr1ba*\* and MZ*oep* + *acvr1ba*\*) represents embryoid velocity and global tissue flow given as arrowhead white vectors (5 μm/min) across the maximum intensity projections of embryoids. The colour bar depicted relative velocity given in μm/min. Embryoids oriented from distal (top) to proximal (bottom). Scale bar: 200 µm.

Tissue flow analysis revealed an increase in the velocity of cells directed towards the proximal part of the embryoid (Fig. 6, bottom rows). In three observed embryoid conditions (wt, wt+*acvr1ba\**, MZ*oep* + *acvr1ba\**) from the shield stage (6 - 6.7 hpf), multiple regions located around the border of the sample showed an increase in velocity. This corresponded to an increase in the budding activity as previously described in Fig. 5. After approximately 6.7 hpf, cells of wt embryoids started to migrate in the proximal direction until 13 hpf. In contrast to wt embryoids cells of wt+*acvr1ba\** and MZ*oep*+*acvr1ba\** undergo different pathways. Vector analysis revealed that at around time point 7.75 hpf wt+*acvr1ba\** and MZ*oep*+*acvr1ba\** embryoid cells at the proximal parts belong to the 1/16 cell injection progeny or in close proximity collectively move towards outer space. At around 9.05 hpf, flow analysis showed that these cells changed their direction towards the inner part of the embryoids. These cells returned to the embryoid by migrating below the cell layers on the surface. At the end of gastrulation (time point 10.5 hpf in the Fig. 6) cells belonging to more external tissue layers migrated in the proximal direction and formed up to two extension tips visible at 13 hpf. In addition, flow analysis showed an increase in maximum velocity in wt+*acvr1ba\** compared to wt and MZ*oep*+*acvr1ba*\* embryoids.

### Expression of *acvr1ba*\* in embryoids induces the expression of the *gsc* fluorescent reporter

To further investigate the robustness of embryoids to the absence of yolk boundaries and signalling molecules from the yolk and YSL, we studied the effect of *acvr1ba*\* on embryoids (Fig. 7A). The injection of mRNA encoding constitutively activated *acvr1ba*\* into one marginal cell (fated to develop into a mesendoderm) at the 16-cell stage of the embryo resulted in Nodal overexpression in the embryo. To identify the effect of *acvr1ba*\* on embryoids, we used the transgenic zebrafish line *Tg(gsc:EGFP),* which specifically labelled the shield and is present in axial mesodermal and endodermal structures at later stages (Schulte-Merker et al., 1994). Peyriéras et al. reported the overexpression of Nodal by injection of *acvr1ba*\* through duplicated expression of the Nodal response gene, *gsc*. In Fig. 7B we show an example of this duplication where one *gsc* reporter expresses in the shield and induces a shield located at 180° relative to the endogenous shield. To trace progeny of constitutively activated *acvr1ba*\* 1/16 cell microinjection, we co-injected mRNA encoding *H2B-mCherry*. We observed the position of *acvr1ba*\*+*H2B-mCherry* (red) and *Tg(gsc:EGFP)* (green) expression for 1.5 hours after embryoids production (time necessary for embryoids to “heal” and express a sufficient amount of the mCherry fluorescent protein) (Figs 7C, S3). At 13 hpf, embryoids were fully elongated. Based on the spatial positions of *acvr1ba*\* and *gsc,* two gene expression categories could be distinguished: “single green and red overlap” (n=11) and “two green spots and one red overlap” (n=3). The presence of these two categories suggested that the constitutively activated *acvr1ba*\* Nodal receptor can upregulate *gsc* reporter expression in embryoids at the same place or in the proximity to the region containing *acvr1ba*\*.

**Fig. 7.**
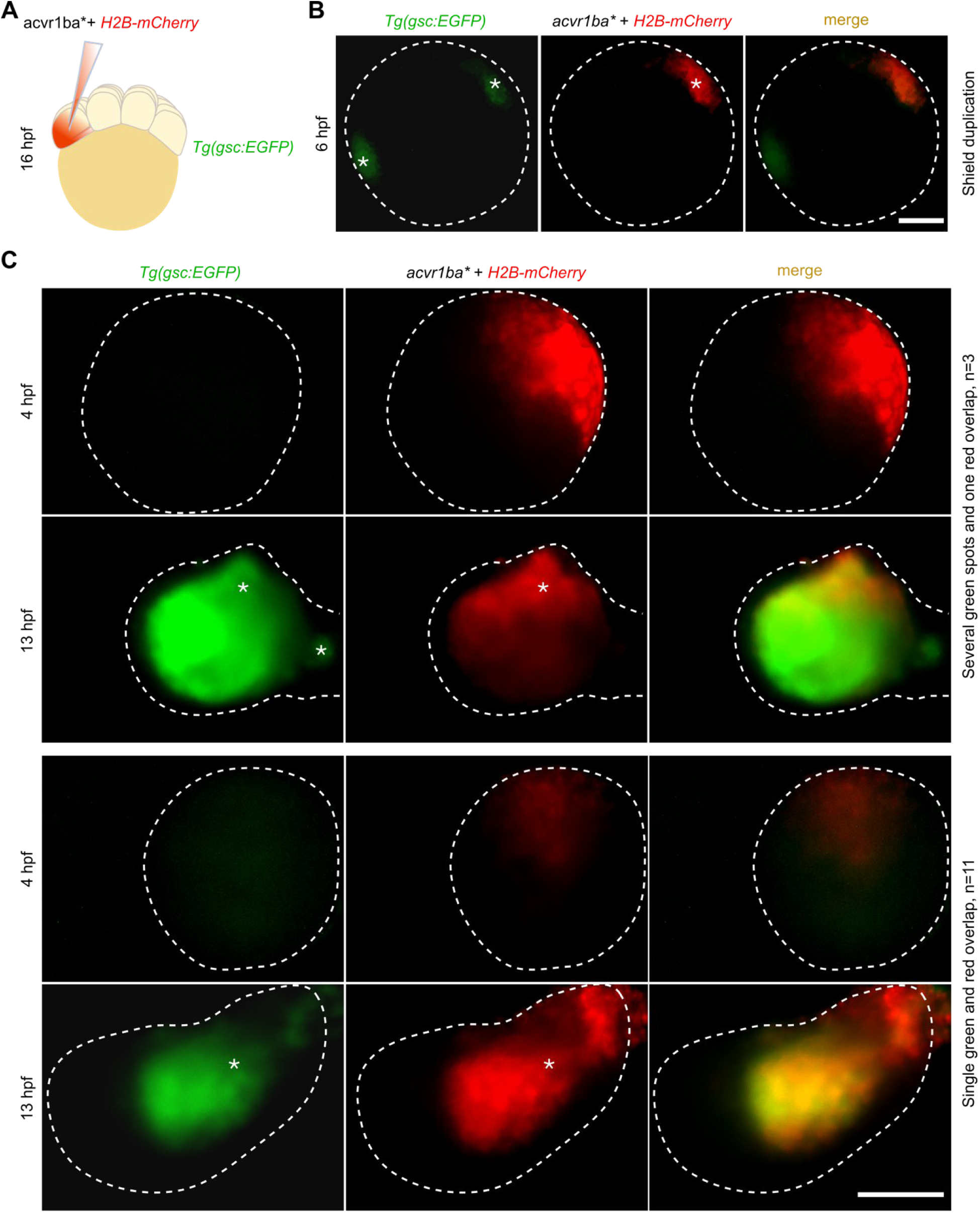
*gsc* reporter expression in embryoids made from 1/16 cell *acvr1ba\** injected embryos. (**A**) Schematic represents coinjection of mRNA encoding *acvr1ba* and H2B-mCherry* into one cell at the 16-cell stage of transgenic embryo *Tg(gsc:EGFP).* (**B**) In embryo injection of acvr1ba* results in the formation of the second shield marked by expression of *gsc* reporter. The image on the left shows *Tg(gsc:EGFP)* (green) expression, the second image represents *acvr1ba* + H2B-mCherry* (red) expression and the third corresponds to merged images of *Tg(gsc:EGFP)* and *acvr1ba* + H2B-mCherry*. (**C**) Two types of spatial distribution of *Tg(gsc:EGFP)* and *acvr1ba* + H2B-mCherry* at 4 hpf and 13 hpf in embryoids: “several green spots and one red overlap” (n=3) and “single green and red overlap” (n=11). Images on the left show *Tg(gsc:EGFP)* expression, the second image corresponds to the progeny of 1 cell injected at 16 cell stage coinjected cell with *acvr1ba*+ H2B-mCherry* and the third image on the right represents merged first and second images of the same row (N=3, n=14). Borders of each fluorescent image are represented with white dashed lines. Spots of *gsc* expression and *acvr1ba* + H2B-mCherry* are denoted with white asterixis. Scale bars: 200 µm.

**Fig. S3.**
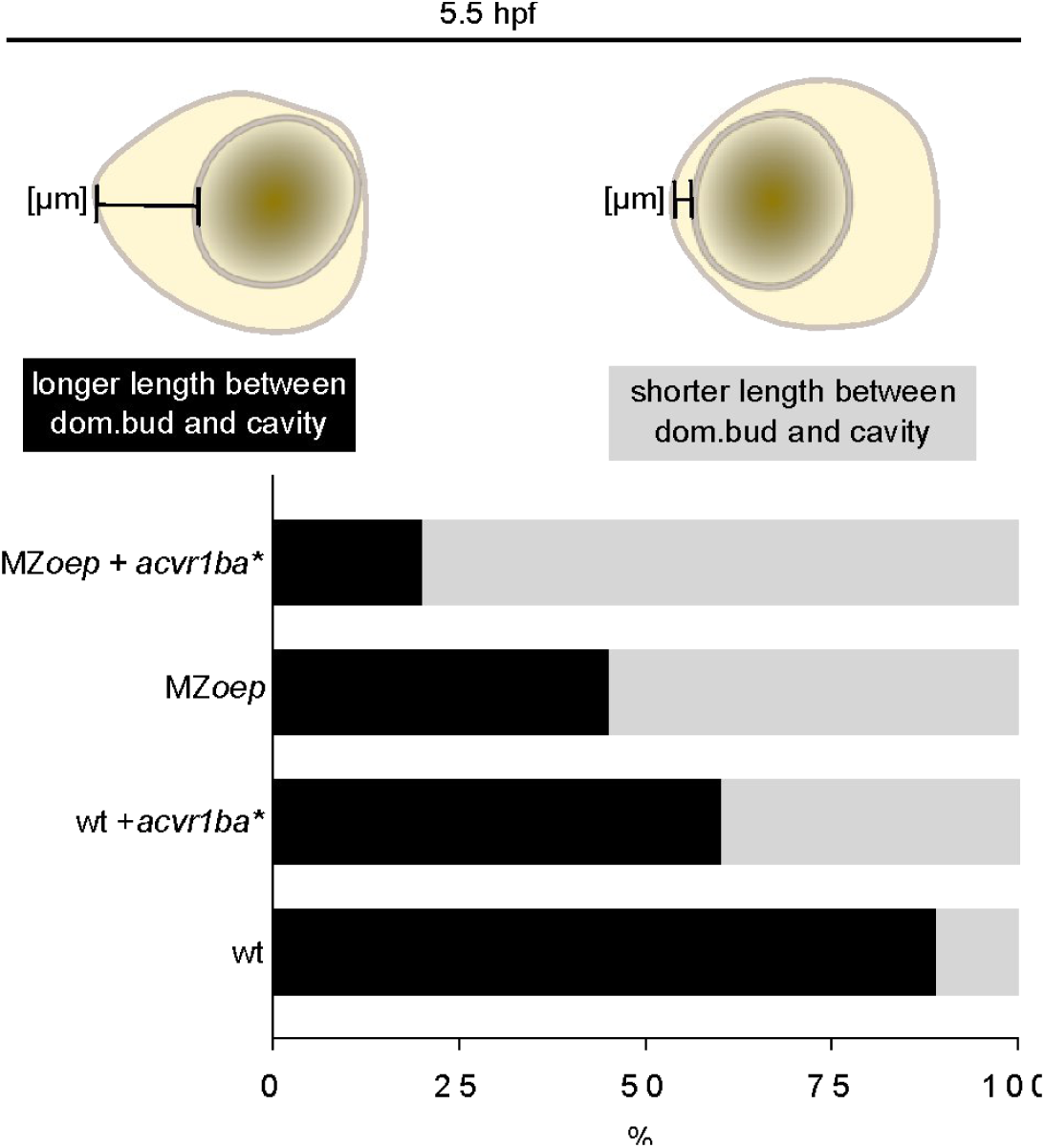
Distance between the elongation pole and the blastocoel like cavity in embryoids depending on Nodal activity. Schematic represents measurement at 5.5 hpf of the length between the edge of the cavity (circular structure) and the highest point of the dominant bud (invagination-like structure on embryoid surface). More than 50% of wt and wt+*acvr1ba\** embryoids have dominant bud most distant from the cavity (wt, N=2 n=9; wt+*acvr1ba*,* N=1 n=5). Contrary to that dominant bud in MZ*oep* and MZ*oep*+*acvr1ba\** embryoids is the most distant in less than 50% of measured samples (MZ*oep*, N=2 n=9; MZ*oep*+*acvr1ba\**, N=1 n=5). Dom., dominant bud.

## Discussion

We addressed cell behaviour during gastrulation without the yolk using the embryoid model system. Moreover, we investigated axis elongation and gastrulation movements as a function of Nodal signalling conditions using embryoids.

Our data showed that selected cell populations obtained from cell lineage tracking indicate impaired convergence; however, extension was intact (Fig. 2). Similar cell behaviour during elongation of presomitic mesoderm was observed by Thomson et al., 2021. They suggested that elongation of the presomitic mesoderm appears in the absence of local cell intercalation by the mechanism referred as compaction-extension. This mechanism has been associated with the reduction of volumetie, higher cell density, and increase in cell volume. Our observation of the presence of extension movement on embryoids, together with previous observations on a similar model (Fulton et al., 2020; Schauer et al., 2020; Williams and Solnica-Krezel, 2019), indicate the high level of conservation of this morphogenetic movement.

We aimed to understand whether yolk deficiency in embryoids disturbed cell behaviour in terms of cell proliferation and mixing. We noted that cells in embryos and embryoids are similar in terms of proliferation rate and mixing (Fig. 3). Our measurement of cell cycle duration supported previous discoveries that cell proliferation is not the only mechanism involved in the first 12 hours of embryoid development (Fulton et al., 2020b; Schauer et al., 2020b; Williams and Solnica-Krezel, 2019). Finally, a similar level of cell proliferation and mixing indicates the robustness of this *in vitro* model to the absence of yolk topology and signalling.

Measurements of the aspect ratio of embryoids consisting of four Nodal signalling conditions (wt, wt+*acvr1ba*\*, MZ*oep* and MZ*oep*+*acvr1ba*\*) provided a better understanding of the involvement of this signalling in the elongation of the body axis (Fig. 4, Movie 3). Wt and wt+*acvr1ba*\* embryoids elongated the most, indicating the involvement of Nodal signalling in this process. The ability to form the second axis extension in wt+*acvr1ba*\* embryoids suggested the significance of Nodal in axis elongation. Evidence that *acvr1ba*\* 1/16 cell injection influences elongation is especially visible from 3D+time data of MZ*oep*+*acvr1ba*\* embryoids, in which cells belong to the progeny of *acvr1ba*\* leading cell movement that forms proximal extension tip (Movie 4). Aspect ratio measurements of MZ*oep*+*acvr1ba*\* embryoids showed higher values than in MZ*oep*. However, elongation occurred in 29.7% of embryoids in MZ*oep* mutants. The elongation of MZ*oep* embryoids confirmed a non-autonomous function of Nodal signalling in this process. In conclusion, modulation of the Nodal pathway in embryoids revealed that Nodal is significant, but not the only factor required for the axis elongation.

We further focused on understanding how modulations in the Nodal pathway affect embryoids morphology (Fig. 5). 2D and 3D live images of embryoids belonging to four Nodal conditions revealed budding activity. Budding occurred from approximately 4.5 hpf until one or two buds took over and formed the extension tip (Fig. 5A). We observed a correlation between embryoids prepared from wt+*acvr1ba*\* embryos and enhanced budding activity reflected by the bud length and number (****p<0.0001) (Fig. 5B,C). In addition, bud length and number (****p<0.0001) with the longest lifetime were identified in wt+*acvr1ba*\* condition (****p<0.0001). The increase in the lifetime of the bud domain in wt+*acvr1ba*\* embryoids indicates greater stability of the Nodal positive loop upon 1/16 cell injection of *acvr1ba*\* (Fig. 6D). Budding has been described during the development of the salivary gland, pancreas, mammary gland, and airway epithelium in the lung (Shih et al., 2013; Spurlin and Nelson, 2017). In these organs, mesenchyme-secreted growth factors belonging to the identical protein superfamily as Nodal (TGF-β) have been reported to be involved in the budding movement (Wang et al., 2021). Our results show the first time the presence of budding activity in zebrafish. However, further research needs to investigate the mechanisms of bud formation and its function.

Previous research has shown that transplanted *acvr1ba*\* naïve cells can induce internalisation in zebrafish. Moreover, this movement is absent in wt embryoids (Schauer et al, 2020). This suggests that *acvr1ba*\* may be involved triggering this movement in the absence of the yolk. Therefore, we explored the possibility to induce internalisation in wt+ *acvr1ba*\* and MZ*oep*+*acvr1ba*\* embryoids. Using 3D+time data, we were able to identify potential internalisation gastrulation movements (Fig. 7). The internalisation-like movements emerged after the shield stage and continued until the bud stage, when cells belonging to the more external layers took over and extended, creating the second extension tip. From 10-13 hpf, both types of embryoids form two extensions. MZ*oep*+*acvr1ba*\* developed shorter extension tips, consistent with the disrupted Nodal signalling pathway in MZ*oep* mutants. In a research article published in 2001, David and Rosa et. al have shown that the activation of Nodal signalling by *acvr1ba*\* in naïve cells can lead to cell internalisation behaviour, which is in agreement with our findings on embryoids. Furthermore, we do not report internalisation-like movements in wt, as it has been recently reported on the same model (Schauer et al, 2020). This supports our hypothesis that 1/16 cell injected with *acvr1ba*\* is responsible for internalisation-like movement of the embryoid.

To further investigate the influence of the Nodal pathway on this model, we focused on its modification upon *acvr1ba*\* 1/16 cell injection. Utilising embryoids prepared from *Tg(gsc:EGFP)* embryos upon *acvr1ba*\* 1/16 cell injection showed expression of two spots of shield and axial mesoderm markers, *gsc*, and even in some instances, second extension tip (Fig. 7); in embryos this behaviour corresponds to the formation of two shields and further the formation of the second body axis (Peyriéras et al., 1998). *Gsc* expression pattern directly indicates a modified Nodal pathway. Considering that the *gsc* expression domain and body axis duplication upon *acvr1ba*\* injection resembles the behaviour in embryos, these results indicate the robustness of embryoid development to the absence of signalling from the yolk.

We showed that zebrafish embryoids are robust to the deficiency of yolk topology and yolk-derived signalling during gastrulation. This work provides a step forward in the integration of molecular and cellular processes underlying gastrulation. Moreover, by adding nutrients to our saline-based culture conditions, this *in vitro* zebrafish system might be a valuable model for further investigations of morphogenesis and differentiation.

## Materials and methods

### Fish lines and husbandry

Zebrafish (*Danio rerio*) experiments and husbandry were all carried out adhering to the rules set by Directive 2010/63/EU of the European Parliament, 2010. Zebrafish handled in E3 medium at 28.5°C (Kimmel et al., 1995).

### Embryoid culture medium

Culture medium containing: 116 mM NaCl, 2.9 mM KCl, 2.4 mM CaCl_2_, 5.0 mM HEPES, pH 7.2, 50 U/ml Penicillin and 0.05 mg/ml Streptomycin (Cat# 15140122, ThermoFischer Scientific).

### Embryoid fixation

Live embryoids were gently taken by glass pipette and transferred in a 3 cm Petri dish filled with 4% PFA in PBS (under the laminar). Embryoids were left for the next 12 hours at 4°C. Next day, embryos were rinsed in PBS 0.1% Tween 20 at least three times for 5 minutes by swirling the plate from time to time. Then, samples were stored at 4°C in PBS 0,1% Tween 2 until used for 3D imaging.

### Embryo microinjection

Respective plasmids encoding *H2B-mCherry*, *EGFP-Hras*, *mCherry-F* and *acvr1ba*\* were linearised with the appropriate enzyme and purified by Phenol-Chloroform extraction. According to the manufacturer’s procedure, mRNA was transcribed using the mMessage mMachine kit for SP6 (Ambion). According to the manufacturer’s procedure, mRNA was then purified with the Nucleospin RNA XS kit (Machery-Nagel). Zebrafish mutant line MZ*oep*^tz57/z57^ was injected at the one-cell stage with 100 ng/µl *H2B-mCherry* and *Hras-EGFP* mRNA. Zebrafish lines *Tg(gsc:EGFP)* SD5 and MZ*oep*^tz57/z57^ mutant, were used for 1/16 cell injection. *Acvr1b\** and *H2B-mCherry* mRNA or *acvr1ba*\* and *mCherry-F* mRNA were injected into zebrafish eggs at the 16-cell stage into one of the 12 marginal blastomeres. The concentration of *acvr1ba*\* was 0.35 ng/µl and 100 ng/µl for *H2B-mCherry* and *mCherry-F*. Zebrafish lines *Tg(act26:mcherry-F)* were used for 64-128 cell stage injection of 12.5 µM SYTOX Green fluorescence dye (Cat# S7020, Invitrogen) in the space of YSL.

### Image acquisition

Embryoids mounted inside the wells (r_1_=600 µm, r_2_=500 µm, h=1000 µm) made by using custom designed embryoid 3D printed mold. Mold was left on the top of 3 cm Petri dish which is filled for 25% with melted 1% agarose (Cat# 50070, BioProducts) diluted in embryoid culture medium. Embryos were dechorionated and mounted into a Teflon ring, inside a 3 cm Petri dish filled with E3 medium. Temperature of 28.5°C maintained with the OKOlab temperature control system (DGT-CO2BX-PLUS, OKOlab, Italy) for the 3D+time imaging and the Minitube HT 50 heating system for microscopes (#12055/0050, Minitube GmbH, Germany) in fluorescence imaging and bright field two-dimensional imaging. 3D+time image acquired using LSM780 Zeiss one-photon and SP5 Leica 2-photon microscopy with 20x/1.0 Zeiss water immersion objective and Olympus 20x/0.95 water immersion objective, respectively. For one-photon microscopy sample excitation continuous lasers were used (λ=488 and λ =543 nm) and two photon microscope uses Spectra-Physics Mai Tai Ti:Sapphire, λ =980 nm and Amplitude, λ=1030 nm femtosecond lasers. Two-dimensional bright field image acquisition was obtained with Leica MZ16 microscope with PLANAPO 1.0 x objective with chosen magnifications of: 1.6x, 2x and 2.5x. Two-dimensional bright field and fluorescent images were acquired using BX61 Olympus microscope with 5x/0.5 Nikon air objective, CoolLED p*E*-4000 illumination system (λ=490 nm and λ=550 nm) and Orca Flash 4.0 Hamamatsu camera for detection. Image acquisition parameters are given in the Tables S2, S3, S4 and S5.

**Fig. S4.**
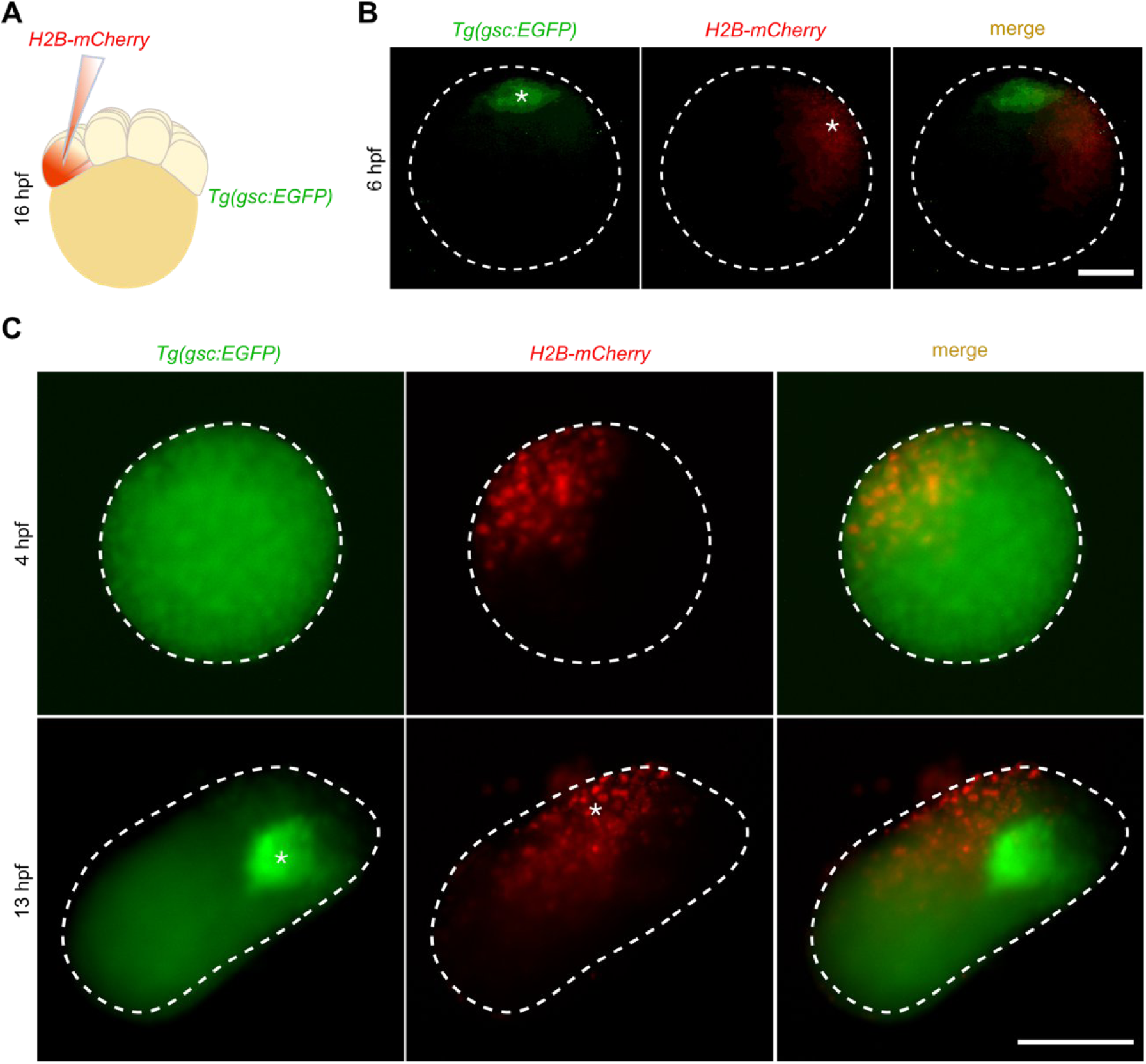
*Gsc* fluorescent reporter expression upon 1/16 cell control injection. Related to. Fig. 7. (**A**) Schematic represents injection of mRNA encoding *H2B-mCherry* into one cell at the 16-cell stage of *Tg(gsc:EGFP)* embryo. (**B**) In embryo (images taken from the animal pole view) progeny of *H2B-mCherry* 1/16 cell injection did not induce formation of second *gsc* reporter region. The first row represents merged images of *Tg(gsc:EGFP)* (green) and *H2B-mCherry* (red) expression, the second row shows *Tg(gsc:EGFP)* expression and the third row corresponds to the progeny of the 1/16 injected cell with H2B-mCherry. (**C**) Spatial distribution of *Tg(gsc:EGFP)* and H2B-mCherry at 4 hpf and 13 hpf. Merged fluorescence images first row, the second row shows *Tg(gsc:EGFP)* and the third row corresponds to the progeny of the 1/16 cell injected with *H2B-mCherry* (N=3, n=6). Borders of each fluorescent image are depicted with white dashed lines. Spots of goosecoid (green) expression and *H2B-mCherry* (red) are denoted with white asterixis. Samples oriented from proximal (top) to distal (bottom). Scale bars: 200 µm.

**Table S2.**
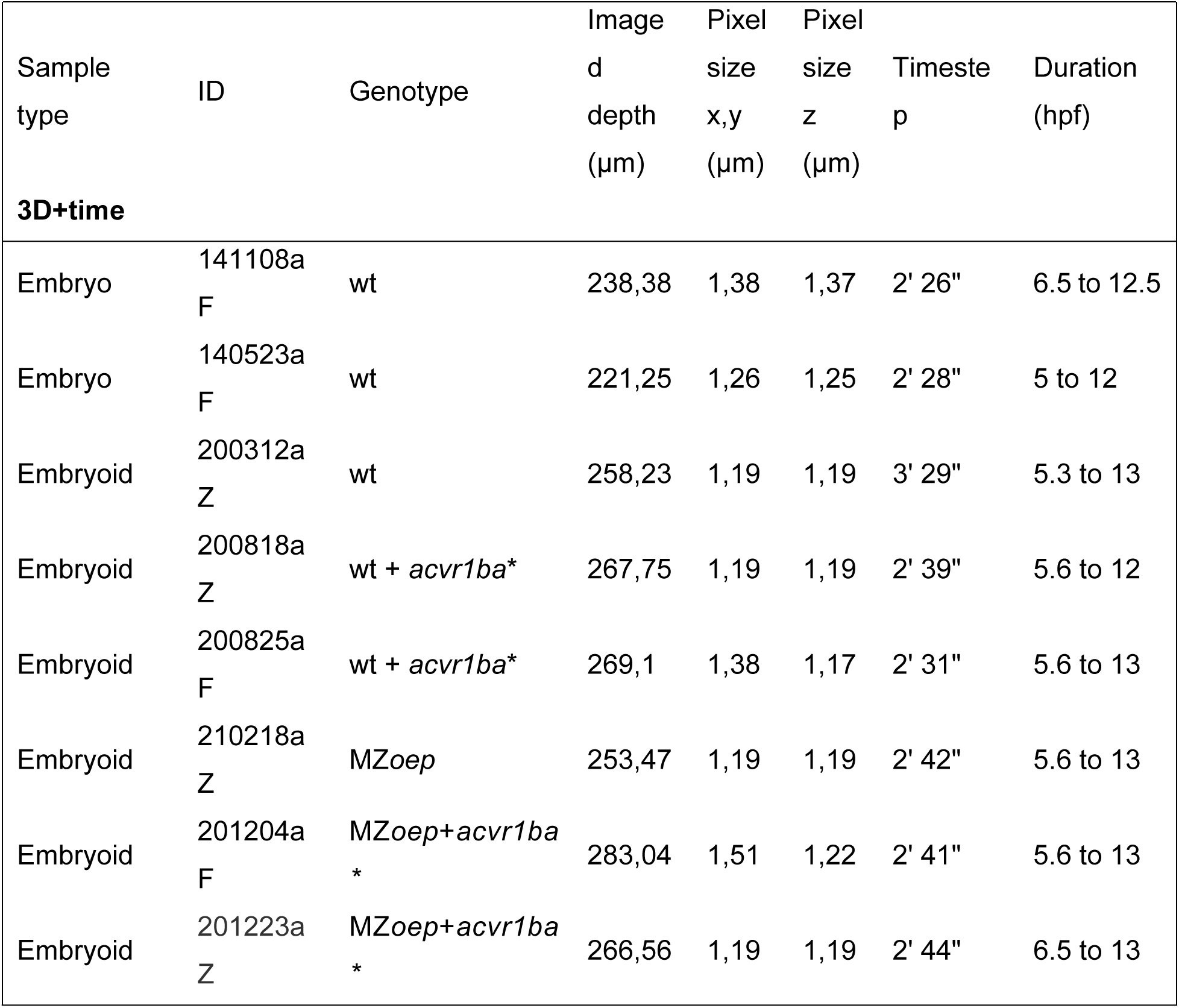
3D+time imaging parameters.

**Table S3.**
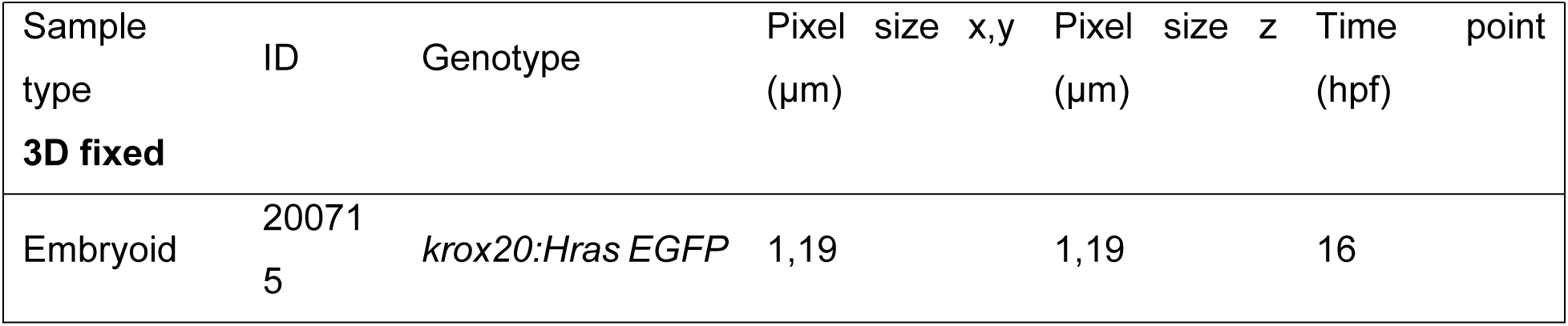
Image acquisition parameters of 3D fixed data.

**Table S4.**
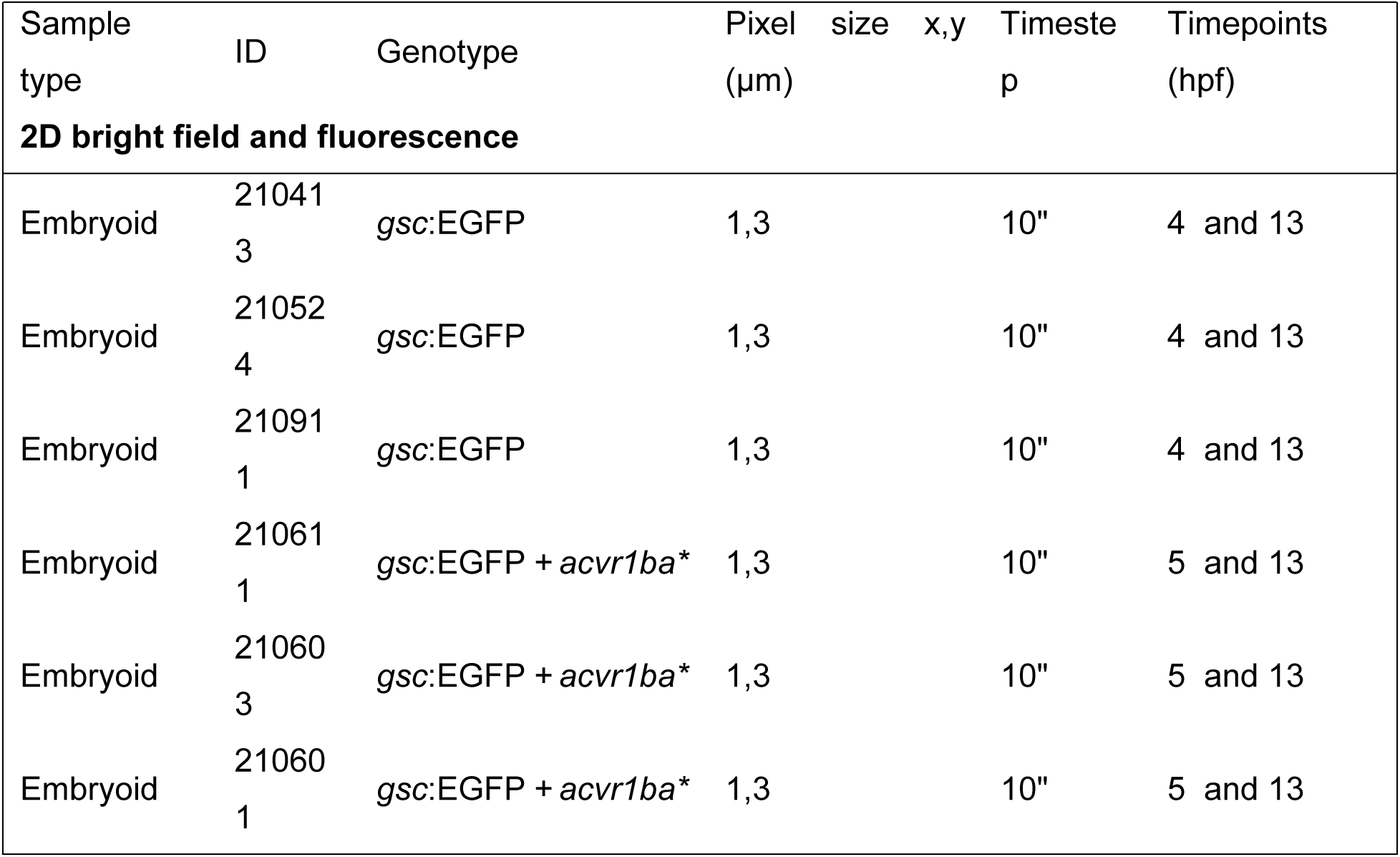
Image acquisition parameters for 2D bright field and fluorescence data.

**Table S5.** Image acquisition parameters for 2D bright field data.

| Sample type | ID | Genotype | Magnification | Pixel size x, y (µm) | Time step | Duration (hpf) |
| --- | --- | --- | --- | --- | --- | --- |
| **2D bright field** |  |  |  |  |  |  |
| Embryoid | 200805 | wt | 1,25 | 4,032 | 1" | 5.5 to 13 |
| Embryoid | 200807 | wt | 1,25 | 4,032 | 1" | 5.5 to 13 |
| Embryoid | 210119 | wt | 1,6 | 3,15 | 1" | 5.5 to 13 |
| Embryoid | 200818 | wt+ *acvr1ba* * | 2 | 2,52 | 1" | 5.5 to 13 |
| Embryoid | 200902 | wt+ *acvr1ba* * | 2,5 | 2,016 | 1" | 5.5 to 13 |
| Embryoid | 200825 | wt+ *acvr1ba* * | 1,6 | 3,15 | 1" | 5.5 to 13 |
| Embryoid | 200807 | wt+ *acvr1ba* * | 1,25 | 4,032 | 1" | 5.5 to 13 |
| Embryoid | 21010<br>5 | wt+acvr1ba* | 1,6 | 3,15 | 1" | 5.5 to 13 |
| Embryoid | 20072<br>3 | MZoep | 1,6 | 3,15 | 1" | 5.5 to 13 |
| Embryoid | 20082<br>0 | MZoep | 1,6 | 3,15 | 1" | 5.5 to 13 |
| Embryoid | 20091<br>7 | MZoep | 1 | 5,04 | 1" | 5.5 to 13 |
| Embryoid | 20110<br>5 | MZoep | 1,6 | 3,15 | 1" | 5.5 to 13 |
| Embryoid | 21021<br>8 | MZoep | 1,6 | 3,15 | 1" | 5.5 to 13 |
| Embryoid | 20120<br>4 | MZoep+acvr1ba<br>* | 1,6 | 3,15 | 1" | 5.5 to 13 |
| Embryoid | 20122<br>3 | MZoep+acvr1ba<br>* | 1,6 | 3,15 | 1" | 5.5 to 13 |
| Embryoid | 21030<br>1 | MZoep+acvr1ba<br>* | 1,6 | 3,15 | 1" | 5.5 to 13 |
| Embryoid | 21031<br>7 | MZoep+acvr1ba<br>* | 1,6 | 3,15 | 1" | 5.5 to 13 |

### Analysis of SYTOX positive domains

SYTOX domain measurement has been done by using Analyse Particles tool that is a part of Fiji software (Schindelin et al., 2012). Percentage of SYTOX Green domain was measured as a ratio between the area of SYTOX Green fluorescence and the area of the embryoid obtained from bright field two-dimensional images. Images used in measurement obtained 30 min after 256-cell stage embryo dissection which corresponds to 3 hpf embryoid. A detailed value can be seen in the Excel sheet.

### Analysis of embryoid buds

Analyse Particles function in Fiji software was used to analyse the morphology of two-dimensional bright field images: number of buds per embryoid, bud length and dominant bud duration. Detailed values are included in the Excel sheet.

### Statistics

GraphPad Prism 9 software was used for statistical analysis and to generate graphs. The statistical test varies between experiments, and they are described in figure legends and the text. To compare two groups unpaired t-test was used and to compare four groups two-way ANOVA test.

### Image processing

Images belonging to the Fig. 7 were corrected for brightness and contrast. Image belonging to the Figs 1, 6 and Movies 4, 5 were filtered using a custom-made algorithm based on deep learning.

### Tissue flow analysis

We used PIVlab implemented in Matlab to analyse the tissue velocity. Interrogation windows of 32×32 pixels and sub-windows of 16×16 pixels with an overlap of 50% were chosen. Images of embryoids and embryos were pre-processed in ImageJ/Fiji. For each time point, we created a maximum z-projection of the z-stack. Drift corrections using the Template Matching plugin, smoothening and threshold (B&W) adjustments were applied. Heatmaps visualising the velocities were smoothened via interpolations.

### Analysis of cell behaviour

Cell lineage trees were reconstructed with a custom image processing workflow that performs nuclei detection and tracking. Data validation, visualisation and quantitative analysis were performed with Mov-IT software (Faure et al., 2016). Mov-IT can extract numerical information about cell division of each cell lineage which was then used to plot mixing and proliferation. For the stripe analysis, automatically detected cells and lineages of selected cell populations were propagated forward in time to investigate convergence and extension movements in embryos and embryoids.

## Acknowledgements

We would like to thank BioEmergences lab members for discussion and comments on experiments, data interpretation and manuscript. We thank Vikas Trivedi for discussion and ideas. We thank Monique Fran and Mageshi Kamaraj for *Tg(krox20:EGFP H-Ras)* zebrafish line. Maxime Comberiati, Manon Mehraz, Fanny Husson and Animal facility members for help with work on zebrafish.

## Author contributions

Conceptualisation, S.J. and N.P.; Methodology, S.J. and N.P.; Investigation, S.J., N.P., T.S. and J.E.; Data analysis, S.J., and J.E.; Visualisation S.J. and T.S.; Supervision, S.J. and N.P.; Writing -original draft, S.J.; Writing -reviewing & editing, S.J., N.P., J.E., and T.S.; Project Administration, N.P. and T.S.; Resources, N.P.; Funding, N.P.

## Data availability

Datasets are available on the website, https://bioemergences.eu/SJovanic/index.php Correspondence and material requests should be addressed to Nadine Peyriéras.

## References

1. Aoki, T. O., Mathieu, J., Saint-Etienne, L., Rebagliati, M. R., Peyriéras, N. and Rosa, F. M. (2002). Regulation of Nodal Signalling and Mesendoderm Formation by TARAM-A, a TGFβ-Related Type I Receptor. Developmental Biology 241, 273–288.

2. Carmany-Rampey, A. and Schier, A. F. (2001). Single-cell internalization during zebrafish gastrulation. Current Biology 11, 1261–1265.

3. Chen, Y. and Schier, A. F. (2001). The zebrafish Nodal signal Squint functions as a morphogen. Nature 411, 607–610.

4. Dougan, S. T., Warga, R. M., Kane, D. A., Schier, A. F. and Talbot, W. S. (2003). The role of the zebrafish *nodal* -related genes *squint* and *cyclops* in patterning of mesendoderm. Development 130, 1837–1851.

5. Duboule, D. (1994). Temporal colinearity and the phylotypic progression: a basis for the stability of a vertebrate Bauplan and the evolution of morphologies through heterochrony. Development 1994, 135–142.

6. Feldman, B., Gates, M. A., Egan, E. S., Dougan, S. T., Rennebeck, G., Sirotkin, H. I., Schier, A. F. and Talbot, W. S. (1998). Zebrafish organizer development and germ-layer formation require nodal-related signals. Nature 395, 181–185.

7. Fulton, T., Trivedi, V., Attardi, A., Anlas, K., Dingare, C., Arias, A. M. and Steventon, B. (2020). Axis Specification in Zebrafish Is Robust to Cell Mixing and Reveals a Regulation of Pattern Formation by Morphogenesis. Current Biology 30, 2984–2994.e3.

8. Gilbert, S. F. and Gilbert, S. F. (2000). Developmental Biology. 6th ed. Sinauer Associates.

9. Gritsman, K., Talbot, W. S. and Schier, A. F. Nodal signaling patterns the organizer. 12.

10. Holtfreter, J. (1944). A study of the mechanics of gastrulation. Journal of Experimental Zoology 95, 171–212.

11. Huch, M., Knoblich, J. A., Lutolf, M. P. and Martinez-Arias, A. (2017). The hope and the hype of organoid research. Development 144, 938–941.

12. Kimmel, C. B., Ballard, W. W., Kimmel, S. R., Ullmann, B. and Schilling, T. F. (1995). Stages of embryonic development of the zebrafish. Developmental Dynamics 203, 253–310.

13. Liu, Z., Woo, S. and Weiner, O. D. (2018). Nodal signaling has dual roles in fate specification and directed migration during germ layer segregation. Development dev.163535.

14. Montero, J.-A., Carvalho, L., Wilsch-Bräuninger, M., Kilian, B., Mustafa, C. and Heisenberg, C.-P. (2005). Shield formation at the onset of zebrafish gastrulation. Development 132, 1187–1198.

15. Ninomiya, H., Elinson, R. P. and Winklbauer, R. (2004). Antero-posterior tissue polarity links mesoderm convergent extension to axial patterning. Nature 430, 364–367.

16. Peyriéras, N., Strähle, U. and Rosa, F. (1998). Conversion of zebrafish blastomeres to an endodermal fate by TGF-β-related signalling. Current Biology 8, 783–788.

17. Schauer, A., Pinheiro, D., Hauschild, R. and Heisenberg, C.-P. (2020). Zebrafish embryonic explants undergo genetically encoded self-assembly. eLife 9, e55190.

18. Schier, A. F., Neuhauss, S. C. F., Helde, K. A., Talbot, W. S. and Driever, W. (1997). The one-eyed pinhead gene functions in mesoderm and endoderm formation in zebrafish and interacts with no tail. Development 124, 327–342.

19. Sepich, D. S., Calmelet, C., Kiskowski, M. and Solnica-Krezel, L. (2005). Initiation of convergence and extension movements of lateral mesoderm during zebrafish gastrulation. Dev. Dyn. 234, 279–292.

20. Soh, G. H., Pomreinke, A. P. and Müller, P. (2020). Integration of Nodal and BMP Signaling by Mutual Signaling Effector Antagonism. Cell Reports 31, 107487.

21. Solnica-Krezel, L. (2005). Conserved Patterns of Cell Movements during Vertebrate Gastrulation. Current Biology 15, R213–R228.

22. Strähle, U., Jesuthasan, S., Blader, P., Garcia-Villalba, P., Hatta, K. and Ingham, P. W. (1997). one-eyed pinhead is required for development of the ventral midline of the zebrafish (Danio rerio) neural tube. Genes Funct 1, 131–148.

23. Turner, D. A., Baillie-Johnson, P. and Martinez Arias, A. (2016). Organoids and the genetically encoded self-assembly of embryonic stem cells. BioEssays 38, 181–191.

24. Wang, S., Matsumoto, K., Lish, S. R., Cartagena-Rivera, A. X. and Yamada, K. M. (2021). Budding epithelial morphogenesis driven by cell-matrix versus cell-cell adhesion. Cell 184, 3702–3716.e30.

25. Williams, M. L. and Solnica-Krezel, L. (2020). Nodal and planar cell polarity signaling cooperate to regulate zebrafish convergence and extension gastrulation movements. eLife 9, e54445.

26. Xu, P., Zhu, G., Wang, Y., Sun, J., Liu, X., Chen, Y.-G. and Meng, A. (2014). Maternal Eomesodermin regulates zygotic nodal gene expression for mesendoderm induction in zebrafish embryos. Journal of Molecular Cell Biology 6, 272–285.

